# Modeling naturalistic face processing in humans with deep convolutional neural networks

**DOI:** 10.1101/2021.11.17.469009

**Authors:** Guo Jiahui, Ma Feilong, Matteo Visconti di Oleggio Castello, Samuel A. Nastase, James V. Haxby, M. Ida Gobbini

## Abstract

Deep convolutional neural networks (DCNNs) trained for face identification can rival and even exceed human-level performance. The ways in which the internal face representations in DCNNs relate to human cognitive representations and brain activity are not well understood. Nearly all previous studies focused on static face image processing with rapid display times and ignored the processing of naturalistic, dynamic information. To address this gap, we developed the largest naturalistic dynamic face stimulus set in human neuroimaging research (700+ naturalistic video clips of unfamiliar faces). We used this novel naturalistic dataset to compare representational geometries estimated from DCNNs, behavioral responses, and brain responses. We found that DCNN representational geometries were consistent across architectures, cognitive representational geometries were consistent across raters in a behavioral arrangement task, and neural representational geometries in face areas were consistent across brains. Representational geometries in late, fully-connected DCNN layers, which are optimized for individuation, were much more weakly correlated with cognitive and neural geometries than were geometries in late-intermediate layers. The late-intermediate face-DCNN layers successfully matched cognitive representational geometries, as measured with a behavioral arrangement task that primarily reflected categorical attributes, and correlated with neural representational geometries in known face-selective topographies. Our study suggests that current DCNNs successfully capture neural cognitive processes for categorical attributes of faces, but less accurately capture individuation and dynamic features.

## Introduction

Deep convolutional neural networks (DCNNs) that are trained for face identification can match or even exceed human-level performance (1–3). Do these models learn internal representations of faces similar to human cognitive and neural representations? Attempts to directly interpret the embedding spaces learned by DCNNs suggest the models may implicitly represent a variety of face features (4). Previous studies reported that representations of objects and faces in deep layers of DCNNs show substantial similarity to neural responses in the ventral temporal cortex of nonhuman primates (5–8). Recent studies reported similar face representations in DCNNs and the human brain (9–13). Nearly all prior studies, however, used static face images with short display times (hundreds of milliseconds). The only study so far (13) that used dynamic naturalistic video clips of faces with longer presentation times (3 s) reported weak correlations between face representations in DCNNs and the brain.

Although face perception processes operate on both still images and videos, the quick processing of static images with rapid display times and the more extended processing of longer dynamic videos may engage different cognitive processes and brain responses. Early processing of still images affords individuation of identity but is only a small part of more extended face processing in naturalistic settings. Recognition of identity appears to be achieved in under 400 ms, but people continue to watch faces intently long after identity is established. The extended processing of naturalistic, dynamic faces may elaborate information that relates inferences of state of mind to social cognitive and semantic context. In support of this view, neural responses to dynamic videos reveal a richer information space that is not evident in responses to static images (14–18). It is currently unclear whether DCNNs capture these additional levels of information about faces.

To test the utility of DCNNs as models of human cognitive and neural representations of dynamic, naturalistic faces, we developed a stimulus set comprising 707 naturalistic 4 s video clips of unfamiliar faces (19). To our knowledge, this face stimulus set, alongside the accompanying fMRI data, is the largest currently available in the neuroimaging literature. Faces in these video clips vary across a broad spectrum of perceived gender, age, ethnicity, head orientations, and expressions, providing a rich sampling of the high-dimensional face space. We analyzed this dynamic face stimulus set in terms of representational geometries produced by DCNNs, by behavioral measures of perceived similarity and categorical attributes, and by fMRI measures of neural responses. To ensure that our results were not dependent on a specific DCNN architecture, we repeated all analyses using three separate face-DCNNs. In a behavioral arrangement task, raters placed thumbnails of face videos in a two-dimensional field according to perceived similarity. In a second behavioral task, raters judged categorical attributes of the face images (gender, ethnicity, age, expression, and head orientation). Instead of limiting our analysis to a few face-selective regions as in previous studies (mainly the occipital and posterior temporal cortices), we compared face representations between DCNNs and cortical responses across the entire face-processing network, including regions in the ventral, dorsal, and anterior core system (20, 21).

Representational geometries derived from DCNN, behavioral, and neural measures were all highly reliable, providing a strong foundation and high noise ceilings for investigating their interrelationships. Correlations between representations in DCNNs and the behavioral arrangement task were high, approaching the noise ceiling. Further analysis with feature ratings showed that representational geometries produced by both DCNNs and the behavioral arrangement task were dominated by categorical face attributes. Even though the final, fully-connected layers of DCNNs are optimized for view-invariant recognition of identity, their correlations with behavioral and neural geometries were markedly weaker than were correlations with late-intermediate layers, suggesting that the human cognitive and neural processes for face individuation are poorly modeled by DCNN processes for face individuation. Correlations of neural representational geometries with DCNN and behavioral representational geometries were significant, albeit low, with a meaningful cortical distribution. The highly-reliable but unexplained variance in neural representational geometries may reflect face information beyond categorical attributes, such as dynamic information that is not captured by the behavioral tasks or by DCNNs, or it may reflect face-identity information that is used by the human brain but not by DCNNs. Overall, our results show that current DCNNs successfully model representations of categorical face attributes, but support our hypothesis that their utility for modeling human cognitive and neural representations of dynamic, naturalistic faces may be limited to this early stage of processing and not extend to information embedded in dynamic information and to human processes for face individuation.

## Results

### Reliable face representations in DCNNs, human behavior, and the brain

To investigate shared information in DCNNs, human behavior, and the human brain, we characterized the representations of 707 naturalistic face video clips with multiple high-performing DCNNs, a behavioral arrangement task of perceived similarity, and fMRI data.

To derive DCNN face representations, we first used InsightFace, a state-of-the-art deep face recognition package (https://github.com/deepinsight/insightface). This package includes face detection (RetinaFace), face alignment, and face recognition (ArcFace) steps (Figure 1A) and is currently the industry standard for face identification. We compared these representations to those in two other face-trained DCNNs (AlexNet and VGG16) and two object-trained DCNNs with the same architecture (AlexNet and VGG16) (22).

**Figure 1.**
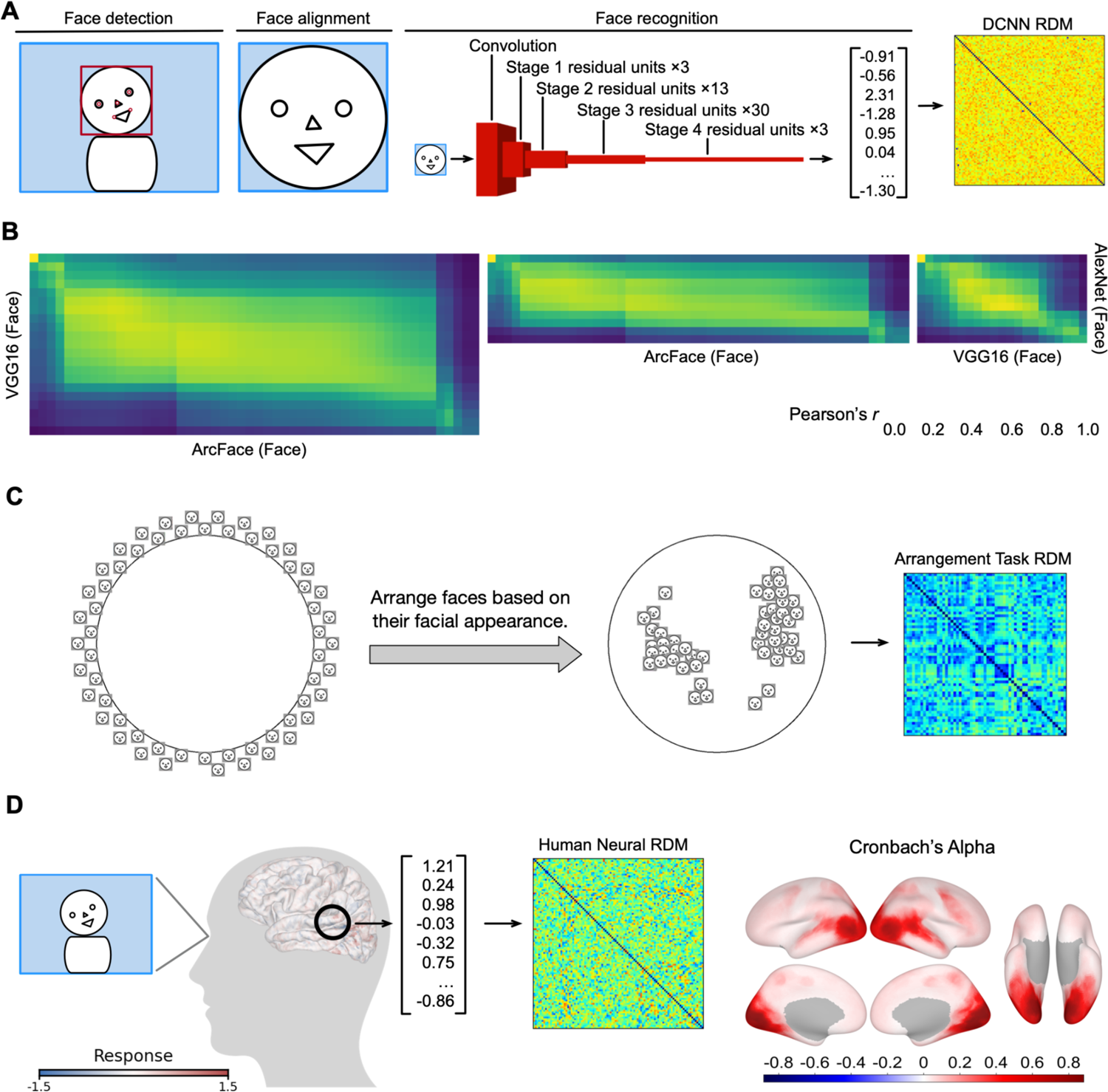
Schematic illustration of representational dissimilarity matrices (RDMs) and reliabilities of deep convolutional neural networks (DCNNs), behavioral performances, and human neural responses. **A.** The DCNN face recognition process comprised three steps. First, the face and its five key landmarks were automatically detected in each frame, and these landmarks were used to create the image of the aligned and cropped face. The cropped face image was then fed into the DCNN as input, and passed through convolutional layers, residual units, and fully-connected layers. The final output was a 512-dimensional embedding vector. Each video clip comprised 120 frames, and the corresponding 120 vectors were averaged to obtain an average embedding vector for each clip. We computed the dissimilarities between the embedding vectors of the 707 face clips to build a 707 × 707 RDM. Note that DCNNs and human subjects were presented with the same naturalistic face videos, and this illustrative example was based on the fully-connected layer of ArcFace. **B.** Correlations in each pair of layers in the three face-DCNN pairs. **C.** In the behavioral arrangement task, MTurk workers organized face stimuli based on facial appearance, and behavioral RDMs were constructed based on the distances between stimulus pairs. Note that this figure is illustrative and not based on real data in the experiment. Mean Cronbach’s alpha across participants was high (0.74). **D.** Human participants watched face video clips in the fMRI scanner, and their brain responses were recorded. For each brain region (searchlight), responses of multiple vertices in the region formed a spatial pattern, and the resulting pattern vector was considered the neural representation of the face clip for that brain region. For each brain region, we computed the dissimilarities between the pattern vectors of the 707 face clips, which formed a 707 × 707 RDM. The surface plot depicts Cronbach’s alphas (noise ceilings) of brain RDMs across all cortical searchlights.

In the behavioral arrangement task, workers on Mechanical Turk (MTurk, https://www.mturk.com/) arranged videos according to perceived face similarities. The stimuli used in single scanning runs (58 or 59 faces) were positioned outside of a circle at the beginning of the task, and MTurk workers were asked to arrange the stimuli inside the circle based on the similarity of facial appearance (Figure 1C). To retain the dynamic aspect of the stimuli, each stimulus would expand when the cursor hovered over its thumbnail and play the 4 s video. The video automatically played once when the cursor hovered the first time, and participants could re-watch the video at any time if they right-clicked the thumbnail.

In the fMRI experiment, human participants underwent scanning while viewing a sequence of 4 s dynamic, naturalistic video stimuli (Figure 1D). Current state-of-the-art fMRI localizers for defining functional face category selectivity use similar dynamic videos of faces in naturalistic settings (23, 24). Brain data from all participants were functionally aligned using hyperalignment based on participants’ brain activity (Figure S1) measured while watching a commercial movie, the Grand Budapest Hotel (25). Hyperalignment aligns brain response patterns in a common high-dimensional information space to capture shared information encoded in idiosyncratic topographies and greatly increases intersubject correlation of local representational geometry (16, 26–30).

Representational dissimilarity matrices (RDMs) were constructed for DCNNs, behavioral similarity arrangements, and neural responses using similar methodologies to characterize pairwise distances between face video clips (see Material and Methods for details). We first assessed the reliability of the information content in the RDMs.

We compared RDMs of different DCNN architectures by calculating correlations between layers of the three face-DCNNs (Figure 1B). Although the three face-DCNNs had different architectures, they shared highly similar representational geometries for faces in our stimulus set, especially in the middle layers (Pearson’s *r* > 0.7). These cross similarities were layer specific, which extended previous results showing layer-specific DCNN representational geometries for objects using object-trained DCNNs (31, 32). Correlations between ArcFace and the other two face-DCNNs in the last few layers and fully-connected layers also were significant (Pearson’s *r* > 0.3) but lower than in the middle layers. We found similarly reliable RDMs for the two object-DCNNs (Figure S3A).

Next, we calculated the noise ceiling for the behavioral arrangement task. Since each behavioral trial showed only face videos from one scanning run (∼60 faces), this task measured RDM similarity across workers for stimuli within scanning runs. The noise ceiling was calculated using Cronbach’s alpha (33), which was computed first across participants within each run and then averaged across runs. A high Cronbach’s alpha means that RDMs from different participants are similar to each other, and that the average RDM has a high signal-to-noise ratio. The mean Cronbach’s alpha value was 0.74, indicating highly similar behavioral arrangements across participants.

We then measured the reliability of neural RDMs across subjects. Noise ceilings were high with maximum values exceeding 0.8 in early visual and 0.7 in posterior face-selective regions (Figure 1D & S12G). Noise ceilings in the anterior face regions were around 0.1–0.4. To further demonstrate the high reliability of neural representations, we conducted additional decoding analyses that revealed high cross-subject face identity decoding accuracies (over 80% accuracy in posterior face areas, Figure S4).

Furthermore, similarities of neural RDMs between areas of the face processing system replicated previous findings describing how face representations change from region to region (20, 34) (Figure S5). Overall, these results showed that meaningful information for faces was reliably encoded in local patterns of fMRI responses in cortical face processing areas.

### Strong correlations between DCNN and human behavioral representations

Representational similarity analysis (RSA) was applied to investigate relationships between DCNN and human behavioral representational geometries. Correlations between face-DCNN RDMs (ArcFace, face-AlexNet, face-VGG16) and behavioral RDMs peaked in late-intermediate layers, and the highest correlations were close to the noise ceiling (Figure 2A). These high correlation values demonstrate that face-DCNN feature spaces for our face video stimuli capture information in human cognitive representations. By contrast, correlations between object-DCNN RDMs and behavioral RDMs were low across all layers (Figure S3B). Taken together, these results show that the training objective (face-identification vs. object-categorization) has a stronger effect than the specific DCNN architecture when modeling human behavioral representations.

**Figure 2.**
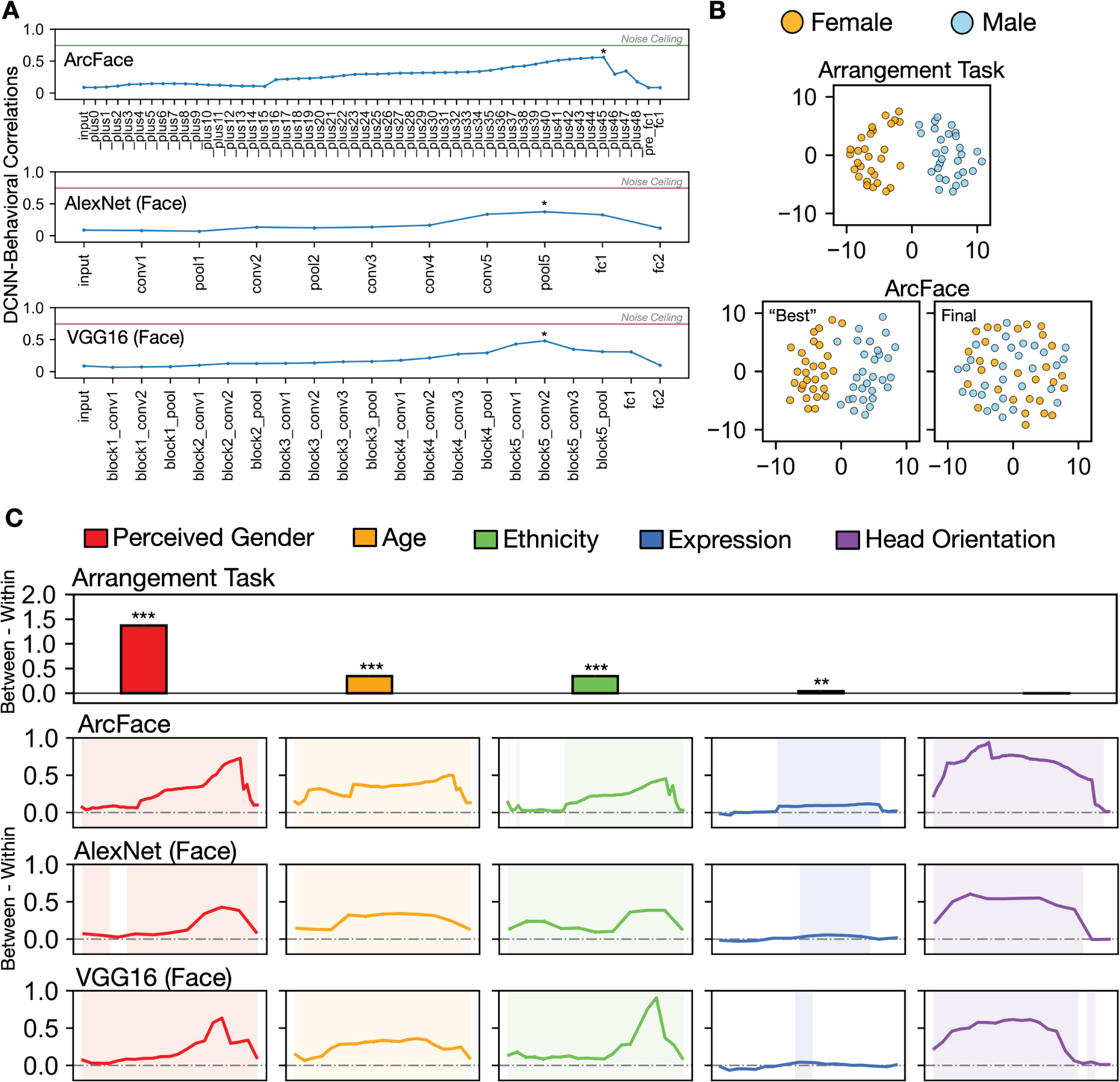
Correlations between DCNN and behavioral, neural and behavioral RDMs. **A.** Mean correlations across participants and runs between the behavioral and DCNN RDMs in each layer in all three face-DCNNs. The star marks the layer that has the highest correlation with the behavioral task in each DCNN. The red horizontal line in each subplot represents the mean noise ceiling of the behavioral arrangement task across runs. **B.** Example MDS plots using RDMs of the same run in the behavioral arrangement task, the “best” layer that showed highest correlation with the behavioral RDM (_plus45) in ArcFace, and the final layer in ArcFace. Each dot is a stimulus. Orange and blue dots indicate perceived females and males, respectively. Behavioral and neural RDMs in this analysis were mean RDMs across participants. **C.** Difference in the between- and within-group distance of perceived gender (red), age (orange), ethnicity (green), expression (blue), and head orientation (purple) in representational geometries of the behavioral arrangement task, and each layer of the three face-trained DCNNs (ArcFace, AlexNet, VGG16). These differences were calculated within each run and then averaged across runs. Shaded layers show significant differences in the between-versus within-group test (*p* < 0.05, permutation test, one-tail). Error bars indicate standard error of the mean estimated by bootstrap resampling the stimuli (the error bars are too small to be visible in some cases). ** *p* < 0.01, *** *p* < 0.001.

To further compare face- and object-DCNNs, we conducted a variance partitioning analysis that quantified how much variance in behavioral representational geometries could be accounted for by face- and object-DCNNs. Results showed that the “best layers” of the face- and object-DCNNs that had the strongest correlations explained 27.5% of the total variance of the behavioral RDMs, due primarily to shared variance with face-DCNNs (27.2% shared variance). By contrast, the final, fully-connected layers accounted for only 5.4% of the total variance of the behavioral RDMs, which was due more to shared variance with object-DCNNs (4.1% shared variance) (Figure S13).

We investigated the nature of the face information that is shared across DCNN and behavioral arrangement RDMs by examining the role of categorical face attributes — perceived gender, age, ethnicity, expression, and head orientation — in face-DCNN and behavioral arrangement RDMs. The contribution of each face feature to the representational geometries was quantified by computing z-scored spatial distances within and between feature groups. Figure 2B shows an example that highlights the rationale of this analysis using multidimensional scaling (MDS) plots. A larger difference of between-group vs. within-group distances corresponds to a clearer division between the feature clusters (e.g., female/male).

Behavioral arrangement RDMs were largely driven by perceived gender, followed by age and ethnicity of faces (Figure 2C). Expression played only a minor role in similarity arrangements, and head orientation played no role. Gender, age, and ethnicity were most strongly reflected in the late-intermediate layers of face-DCNNs (Figure 2C) but only weakly reflected in object-DCNNs (Figure S3D). Head orientation, by contrast, was more strongly reflected in early intermediate layers of face-DCNNs. Thus, the reliable representational geometries in late-intermediate layers of face-DCNNs carry information about cognitive representations that reflect major face categorical attributes. Importantly, representations of these categorical attributes strongly contribute to the similarities between face-DCNNs and the behavioral clustering of perceived similarity.

### Correlations of neural representations with DCNN and cognitive representations

We first correlated neural RDMs in all cortical searchlights with the ArcFace RDMs in each layer. Results showed that representational geometries were more similar in regions extending from the early visual cortex to other regions in the occipital lobe, in the ventral temporal cortex, along the superior temporal sulcus, and in higher-level regions in the frontal lobe for all ArcFace layers (see example maps of the late-intermediate layer _plus45 and the fully-connected layer fc1 in Figure 3A & B). These regions largely correspond to the previously reported human face processing system consisting of multiple face-selective regions (20, 21, 33, 35, 36).

**Figure 3.**
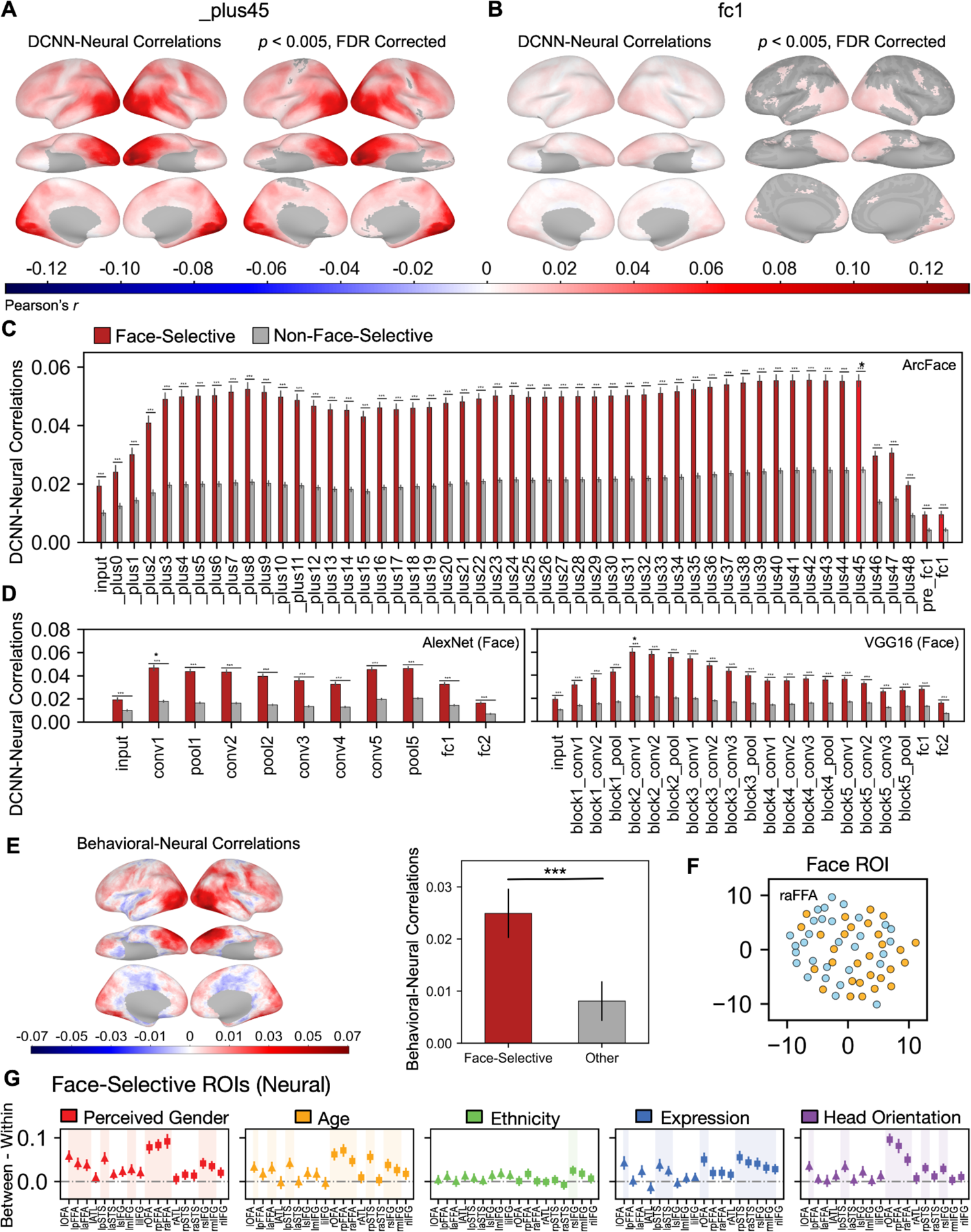
Correlations between DCNN and neural RDMs. **A & B.** The DCNN-neural correlations across all cortical searchlights using RDMs in layer _plus45 (output of the last stage 3 residual unit) and fc1 (the last fully-connected layer). Both layers are highlighted in panel C. Correlations in the visual cortex, ventral temporal cortex, STS, and frontal regions were statistically significant for both layers (controlling FDR at *p* < .005, permutation test). **C.** Average correlations for face-selective regions (defined by a dynamic localizer, faces vs. objects, *t* > 5) and non-face-selective regions (*t* <= 5) are plotted as red and gray bars respectively for each layer. The error bar length stands for one standard error of the mean estimated by bootstrap resampling of stimuli. Significance of the difference between the two bars was assessed via a permutation test randomizing stimulus labels. Layer _plus45 had the largest correlation with neural RDMs among all layers. **D.** Average correlations for face-selective regions and non-face-selective regions for each layer in the two face-DCNNs. Regions, significance, and the color code were defined the same as in Panel C. Stars indicate the layers that had the largest correlations. **E.** The neural-behavioral correlation values in the cortex and the mean behavioral-neural correlations in face-selective (red) and non-face-selective (gray) areas. The error bars indicate standard error of the mean estimated by bootstrap resampling the stimuli. Significance of the difference between the two bars was estimated using a permutation test randomizing the stimulus labels. **F.** Example MDS plots using RDMs for the same run in the right aFFA. Each dot is a stimulus. Orange and blue dots indicate perceived females and males, respectively, the same as Figure 2 panel B. Neural RDMs in this analysis were mean RDMs across participants. **G.** Difference in the between- and within-group distance of perceived gender (red), age (orange), ethnicity (green), expression (blue), and head orientation (purple) in representational geometries of each face-selective ROI (bilateral OFA, pFFA, aFFA, ATL, pSTS, aSTS, sIFG, mIFG, iIFG). These differences were calculated within each run and then averaged across runs. Shaded ROIs show significant differences in the between-versus within-group test (*p* < 0.05, permutation test, one-tail). Significance of the difference was estimated based on a random permutation test randomizing the stimulus labels. Error bars represent one standard error of the mean estimated by bootstrap resampling stimuli. Left triangles are nine face-selective ROIs in the left hemisphere, and right squares are face-selective ROIs in the right hemisphere. *** *p* < 0.001.

We independently defined face-selective regions using a dynamic face localizer (face vs. objects; Figure S6) (24, 33) and calculated the mean correlations for face-selective and non-face-selective regions in each layer. We found that neural RDMs in face-selective regions were best modeled by the late-intermediate ArcFace layers. Correlations dropped drastically after layer _plus45 and reached their lowest values in the final fully-connected layers (Figure 3C). Although correlations in the face-selective regions were significantly higher than the non-face-selective regions in both the peak intermediate layer and the final fully-connected layer, correlations with the peak intermediate layer were more than five times stronger than with the final fully-connected layer across face-selective regions.

To test whether a specific DCNN architecture had a significant effect on the similarities between DCNN and human neural representations, we performed a similar analysis using two other face-DCNNs (AlexNet and VGG16) and found similar results across layers (Figure 3D). Following the same analysis we used for face-DCNNs, we calculated correlations between RDMs in each layer of the two object-DCNNs and neural representations in each searchlight across the cortical sheet. Similarly, mean correlation coefficients were calculated for the face-selective and non-face-selective areas. Correlating object-DCNN and neural RDMs generated comparable correlations to those between face-DCNN and neural RDMs (Figure S3C), indicating that DCNN-neural RDM correlations were not influenced by the training objective of the DCNNs (face-identification vs. object-categorization).

For all DCNNs, intermediate layers provided a markedly better model for the neural representations than final fully-connected layers. This finding is consistent with previous work examining face representation in the brain and DCNNs (5, 10). Interestingly, however, for all DCNNs, correlations between any layer and neural representations accounted for only a fraction of the meaningful variance (all Pearson’s *r* < 0.1). The correlation values were especially low compared to the reliable noise ceilings of neural representations (Figure S7 & S8, the noise ceiling was ∼ 0.4 on average for face-selective areas, with some areas exceeding 0.7, and the DCNN-neural correlations were always < 0.1). Additionally, no meaningful mapping was evident between the layer structure of face-DCNNs and the sequence of face-selective areas in the human neural system for face representations (Figure S12), suggesting that the sequence of representational geometries in the face-DCNN layers differs from the progression of representational geometries along the neural face pathways (20, 34, 37, 38). The DCNN-neural correlations in each individual face-selective ROI for each layer in both face- and object-DCNNs also showed that none of the ROIs had correlations that approached the noise ceiling (Pearson’s *r* < 0.1 in all cases). An additional variance partitioning analysis revealed that the variance in neural RDMs is minimally explained by DCNNs, behavioral models, or by a combination of the two (Figure S13).

We conducted an additional analysis to investigate whether the low correlations were due to RSA’s inherent assumption of equal weights or scales for all features comprising the two RDMs (39–42) (see Material and Methods for details). This additional analysis generated similar results as using classic RSA, excluding the possibility that the low correlations are due to the particular assumption of RSA (Figure S9). To examine whether more distributed brain activity patterns might lead to a better match between the DCNN and neural representations, we repeated this analysis with larger searchlight sizes (15 mm and 20 mm radius). Larger searchlight sizes only slightly improved the correlations, and the overall results remained weak (less than 2% variance explained, Figure S10 & S11).

Correlations between the behavioral arrangement and neural representations in the searchlight analysis consistently showed face-selective areas had significantly higher correlations than other areas (Figure 3E, permutation test, *p* < 0.001). However, the actual correlation values remained small (*r* < 0.1), suggesting that a major difference existed between clustering according to facial similarity and the extended processing of dynamic faces.

The strength of categorical face attributes in neural representational geometries showed a distribution across face-selective ROIs (Figure 3G) that was consistent with known specialization. Face ROIs in the right hemisphere represented these features more prominently than did face ROIs in the left hemisphere. Identity-related invariant categorical face features, such as perceived gender and age, were significantly represented in bilateral face areas in the ventral temporal cortex (OFA, pFFA, aFFA), as well as in the pSTS and right IFG (Figure 3F & G). Expression was significantly represented in face areas in the OFA, pSTS, aSTS, and right IFG, but not the FFA. Head orientation was significantly represented in the OFA, right pFFA and mFFA, and the pSTS. Although neural representations contained significant information for all categorical attributes, this categorical information was more weakly represented than in behavioral and DCNN representations in intermediate layers.

## Discussion

State-of-the-art DCNNs trained to perform face identification tasks have drawn considerable attention from researchers investigating face processing in humans and nonhuman primates. These artificial networks can identify faces at levels of accuracy that match or exceed human performance. Previous neuroscientific studies mainly focused on face representations that are produced by rapid processing of still images. Only one previous study (13) investigated the representations produced by more extended processing of naturalistic, dynamic faces. Here, we investigated the extent to which DCNNs can model real-world face processing by comparing representational geometries produced by DCNNs to representational geometries produced by brain and behavioral responses to a large, varied set of naturalistic face videos.

Our results showed that DCNN, behavioral, and neural representational geometries were stable and information-rich. Face-DCNNs and behavioral representations of perceived similarities captured shared information about face categorical attributes. This face categorical knowledge was strongly represented in late-intermediate layers of face-DCNNs. By contrast, the final fully-connected layers, which are optimized for face identification, did not carry much categorical information. Brain responses to the face videos had representational geometries that were highly reliable across participants, and reliabilities were highest in face-selective cortical areas. Information in the neural representational geometries was significantly correlated with the information in DCNN and behavioral geometries, albeit weakly. The DCNN-neural correlations had a meaningful cortical distribution, following the full distributed face system in occipital, temporal, and frontal cortices, and were much stronger for late-intermediate layers than for the final fully-connected layers. Correlations between DCNN representational geometries and the other two (cognitive and neural) geometries were much stronger for late-intermediate layers too, suggesting that the optimization in the final fully-connected DCNN layers for recognition of identity is a poor model for how human cognitive and neural systems individuate faces.

The maximal local correlations between face-DCNN and neural representations show that at most only 3% of the meaningful variance is shared. It is unclear what information in the highly reliable neural representational geometries is unaccounted for by the face-DCNNs. We focus here on two domains of information in dynamic videos that may play a large role in the variance that remains to be explained: information in facial movement and information derived from other cognitive processes that enrich face representations, such as social inferences, memory, and attention.

In comparison to the rapid processing of briefly presented static stimuli, extended processing of dynamic stimuli dramatically alters the neural response to faces both in terms of tuning profiles and representational geometry. Response tuning to static, well-controlled stimuli in face patches is dominated by the presence or absence of faces or their static structural features (37), but tuning for dynamic face stimuli is dominated by biological motion (14, 18, 43–45). In addition, dynamic faces are superior to static faces for the localization of face-selective areas (23, 24, 33, 46), indicating that they better or more fully engage face-related neural processes. In a similar vein, representational geometry in the ventral temporal cortex for static images of animals is dominated by animal category, but representational geometry for videos of naturally behaving animals is dominated by animal behavior.

Although animal category plays a significant role, it is dwarfed by the representation of behavior, which accounts for 2.5 times more variance (16, 30). Based on results cited here, the uniquely dynamic information in face videos may account for over two-thirds of the variance in neural representational geometries. Further research is needed to precisely characterize how these dynamics change the geometry of face representation. While the processing of static face stimuli may appear to be well-modeled by current DCNNs, extended processing of naturalistic stimuli may reveal the deficiency of such models and help focus our attention on how best to improve them.

Temporally extended face processing with dynamic videos may recruit a variety of cognitive features. People automatically make inferences about novel faces — trustworthiness, competence, attractiveness — that can distort representational geometry (47, 48). Person knowledge plays a large role in the representation of familiar faces (20, 21, 49–52). Familiarity is also known to distort face representations (53, 54), and similarity of novel faces to familiar faces may influence perception and attribution. Faces also play a role in directing attention (36, 55–57), and attention has a large effect on neural responses to faces (58–60) that can be influenced by factors such as trait inferences, familiarity, and memory. Teasing apart the roles played by these different social and cognitive factors on human face representational geometry requires further research. Similarly, developing machine vision systems that incorporate dynamic and social features (expression, eye gaze, mouth movements, etc.) may enhance their power and utility for human-machine interaction.

Behavioral performance in the arrangement task was dominated by major categorical face attributes of perceived gender, age, and ethnicity. These categorical variables play little role in individuation of face identity. In cognitive models of face perception, such categorical judgments precede processing for individuation (61). Many patients with prosopagnosia, who have impaired recognition of face identity, can still judge categorical attributes such as perceived gender, age, expression, and gaze direction (62–64). Thus the categorical face information that is captured by late-intermediate face-DCNN layers and is correlated with cognitive task performance is weakly related to individuation. Processes for face individuation in late, fully-connected layers are powerful but appear unrelated to human cognitive and neural processes.

To summarize the results, Figure 4 illustrates our interpretation of the different components that played a role in the correlations of the DCNN, behavioral, and human neural RDMs. This figure proposes that the high correlations between behavioral RDMs and the RDMs of the intermediate layers of face-DCNNs were mainly driven by the shared categorical information in both types of RDMs. The low correlations with deep layers were due to little face individuation information in the behavioral RDMs as well as little categorical feature information in the RDMs of deep layers. On the other hand, the neural RDMs in the face-processing system contained all four kinds of information – categorical information, face individuation (e.g., Figure S4), dynamic information (dynamic faces are superior to static faces for the localization of face-selective areas), and information from other cognitive processes (e.g., social inference, memory, attention). However, because categorical information contributed the least to the neural RDMs, the shared information between behavioral and neural RDMs was limited. This low contribution of categorical information in neural RDMs can also explain the low correlations between neural RDMs and face-DCNNs in intermediate layers. Finally, the type of information used for face identification in the late, fully-connected DCNN layers and in the human face processing system appear to be quite different. Dynamic information or information from other cognitive processes is essential for the human face processing system (23, 24, 47, 48), while this type of information was not encoded by the face-DCNNs. These differences likely contributed to the low correlations between DCNN RDMs and neural RDMs.

**Figure 4.**
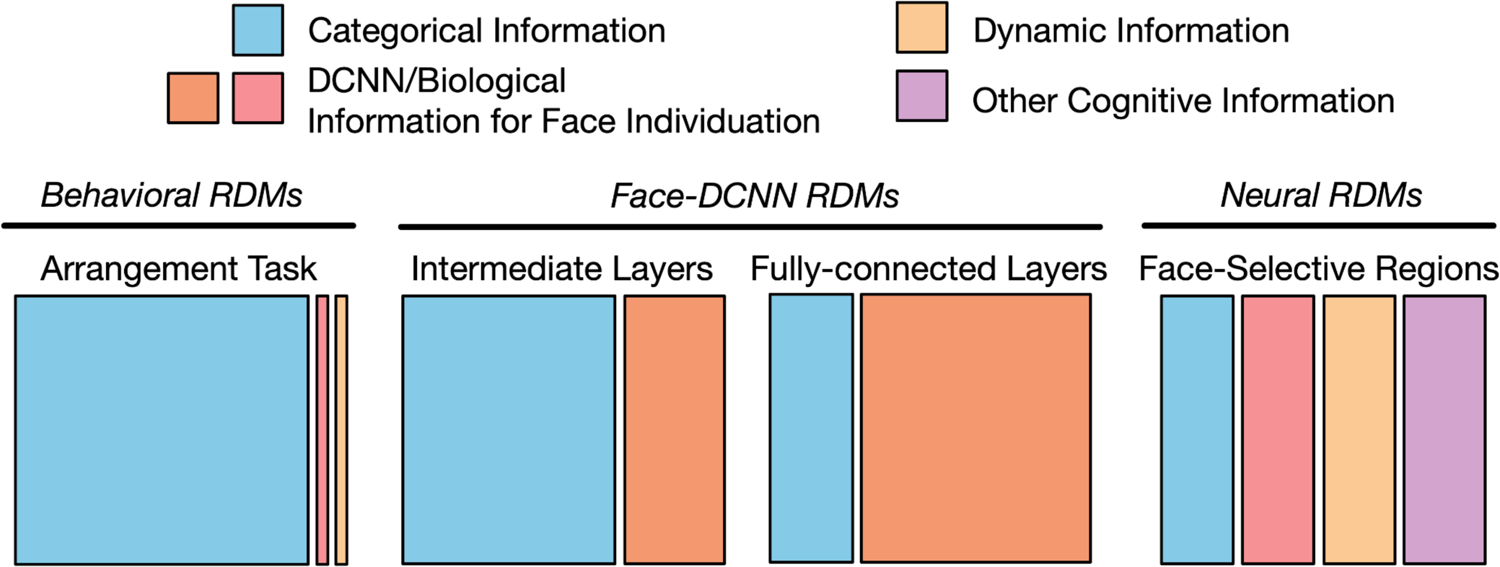
Schematic illustration of hypothesized components of information in RDMs. In RDMs for the behavioral arrangement task and RDMs for the intermediate layers of face-DCNNs, categorical information (blue) played a major role. Most categorical information was factored out in the fully-connected layers of face-DCNNs, and the information for face individuation (orange) became the most prominent part. Neural RDMs in face-selective regions contained similarly weak categorical information as in the final fully-connected layers. Successful individuation of faces may only rely on part of facial features, and the information for face individuation in neural RDMs (red) may be different from that in face-DCNNs. Different from face-DCNNs, neural RDMs also contained dynamic (yellow) and cognitive information (purple) that plays an essential role in human face processing. Note that an extra component existed in all RDMs that stood for the unexplained variance and it was omitted for cleaner display.

Figure 4 suggests a framework for explaining the difference between our results and the results from previous studies that compared DCNNs to brain responses in rapid static face processing tasks. Because dynamic information plays a role in the geometry of brain representations (16, 17, 45), static images could generate higher correlation values between brain responses and DCNNs that do not use motion information (10, 13, 65). Similarly, studies that used stimuli spanning superordinate categories (e.g., with multiple visual categories (12, 42)) would bias representations towards categorical information, reducing the contribution of information that is needed for within-class individuation such as face identification.

Although face-DCNNs are trained on an exceptionally large set of face images, face-DCNNs are optimized to encode these faces according to a very specific objective function: face identification. Face identification, however, is only one aspect of face processing in humans, which is flexible, highly contextualized, and ultimately supports social interaction. Building a representation of the uniqueness of the identity of a face takes a few hundred milliseconds (66), but is followed by sustained processing of a dynamic face in naturalistic viewing for gleaning other information for social cognition — changes of expression gaze, and head-orientation; speech-related mouth movements; inferences of intentions, social rank, social affiliation, reliability, and more. The human system for face perception is serving all of these goals during naturalistic viewing, and processes for face identification, besides playing only a small part that is finished quickly at the onset, may also be integrated with other functions in such a way that identification cannot be simply dissociated as a modular process. Perhaps in the future, artificial neural networks trained with more ecological objective functions (65, 67–69), requiring not just face recognition, but extending to facial dynamics, attention, memory, social context, and social judgments, will learn face representations that afford a more ecologically-valid model that better captures the representations of the face processing system in humans.

## Material and Methods

### Participants

Twenty-one participants (mean age 27.3 years, range 22–31, 11 reported female) participated in the fMRI study. All participants had normal hearing and normal or corrected-to-normal vision, and no known history of neurological illness. The study was approved by the Dartmouth Committee for the Protection of Human Subjects.

### Experimental Design

The Grand Budapest Hotel and localizer data were also used in prior work by Jiahui and colleagues (33). The MRI data acquisition parameters, preprocessing, and data analysis methods involving these two data sets are the same as in the previous publication.

### The Grand Budapest Hotel

The full-length Grand Budapest Hotel movie was divided into six parts. Parts were divided at scene changes to keep the narrative of the movie intact. Participants watched the first part of the movie (∼45 min) outside the scanner. Immediately thereafter, participants watched the remaining five parts of the movie in the scanner (∼50 min, each part lasting 9–13 min) with audio. These data were curated and made publicly available for research use (25).

### Hyperface

Video clips (707 clips, 4 s each) of individuals behaving naturally were created. The video clips were downloaded from YouTube and mostly comprised different people talking in interviews. Individuals in the clips varied widely in their identity, age, ethnicity, perceived gender, and head orientation. Audio channels were removed from the clips and the clips were cropped to remove unrelated text. The video clips were divided into 12 blocks (∼59 clips per block) to match the 12 scanning runs and block order was counterbalanced across participants. In each run, participants were asked to watch the video clips (without fixation), shown continuously. After all clips in a run were shown, participants were tested with a brief four-trial memory check where they were asked to report whether a test clip was novel or was presented in the current run. Feedback was provided at the end of each run. Data from the memory test was removed from all analyses.

### Dynamic Localizer

Participants watched 3 s dynamic clips of faces, bodies, scenes, objects, and scrambled objects (24). The clips were presented continuously in 18 s blocks of each category, without blank periods between blocks. The blocks followed this order: an 18 s fixation period, five blocks of different categories (each lasting 18 s) in random order, an 18 s fixation period, five blocks of the categories in reversed order, and a final 18 s fixation period. Participants were required to press a button whenever they saw a repetition of a clip (five total in each run, one for each category). Four 234 s runs were collected for a total duration of 15:44.

### Behavioral Arrangement Task

An independent group of 39 Amazon Mechanical Turk (MTurk) workers performed this task. Stimuli in a scanning run (59 stimuli for run 1-11 and 58 stimuli for run 12) were displayed as thumbnails outside a white circle on a gray background. When a trial began, the stimuli were arranged in randomized equidistant positions around the circumference of the circle. The first mouse hover triggered a larger and dynamic display of the video clip of that stimulus, and MTurk workers were able to rewatch the video by right clicking the mouse button. MTurk workers were instructed to arrange the thumbnails within the circle based on the similarity of the face appearance. To ensure a reasonable time for each participant to complete the experiment, we asked each of them to perform three trials randomly selected from the total 12 trials. At least 10 different individuals completed each trial.

### Behavioral Rating Task

Another independent group of 121 Amazon Mechanical Turk workers participated in the behavioral rating task. In each trial of the task, participants watched the video clip of a stimulus and rated the stimulus on five features: perceived gender (M/F), age (0-10, 11-20, 21-30, 31-40, 41-50, 51-60, 61-70, 70+), ethnicity (White, Black or African American, Asian, Indian, Hispanic or Latino, Other), expression (Neutral, Happiness, Surprise, Anger, Disgust, Sadness, Fear), and overall head orientation (Mostly Left, Mostly Center, Mostly Right). All 707 stimuli clips were divided into 15 independent experiment sessions (about 47 clips in each session), and each participant was assigned to one session to ensure the experiment could be completed in a reasonable amount of time. At least eight different individuals performed each session, and the final rating of each clip was the one that the most workers agreed on.

### MRI Data Acquisition

All data were acquired using a 3 T Siemens Magnetom Prisma MRI scanner with a 32-channel head coil at the Dartmouth Brain Imaging Center. CaseForge headcases were used to minimize head motion. BOLD images were acquired in an interleaved fashion using gradient-echo echo-planar imaging with pre-scan normalization, fat suppression, multiband (i.e., simultaneous multi-slice; SMS) acceleration factor of 4 (using blipped CAIPIRINHA), and no in-plane acceleration (i.e., GRAPPA acceleration factor of one): TR/TE = 1000/33 ms, flip angle = 59°, resolution = 2.5 mm^3^ isotropic voxels, matrix size = 96 x 96, FoV = 240 x 240 mm, 52 axial slices with full brain coverage and no gap, anterior–posterior phase encoding. At the beginning of each run, three dummy scans were acquired to allow for signal stabilization. The T1-weighted structural scan was acquired using a high-resolution single-shot MPRAGE sequence with an in-plane acceleration factor of 2 using GRAPPA: TR/TE/TI = 2300/2.32/933 ms, flip angle = 8°, resolution = 0.9375 x 0.9375 x 0.9 mm voxels, matrix size = 256 x 256, FoV = 240 x 240 x 172.8 mm, 192 sagittal slices, ascending acquisition, anterior–posterior phase encoding, no fat suppression, and with 5 min 21 s total acquisition time. A T2-weighted structural scan was acquired with an in-plane acceleration factor of 2 using GRAPPA: TR/TE = 3200/563 ms, flip angle = 120°, resolution = 0.9375 x 0.9375 x 0.9 mm voxels, matrix size = 256 x 256, FoV = 240 x 240 x 172.8 mm, 192 sagittal slices, ascending acquisition, anterior–posterior phase encoding, no fat suppression, and lasted for 3 min 21 s. At the beginning of each session (The Grand Budapest Hotel, the Hyperface stimulus, and the localizer task), a fieldmap scan was collected for distortion correction.

### DCNN Models

We used five DCNN models in our analysis: three DCNNs trained for face recognition and two DCNNs trained for object recognition. These DCNNs cover a wide range of commonly used “classic” and state-of-the-art DCNN architectures, including AlexNet (70), VGG16 (71), and ResNet100 (72).

### DCNN Models Trained for Face Recognition

All 3 Face DCNNs were trained using the MS-Celeb-1M dataset (73), one of the largest publicly available datasets for face recognition. The training dataset contains approximately 10 million images sampled from 100,000 top celebrity identities from a knowledge base comprising 1 million celebrities. We used a curated version of the dataset as provided by the InsightFace package, which contains 85,742 identities and 5.8 million aligned face images with 112 × 112 resolution. These images were divided into 45,490 batches with 128 images each during training.

The main architecture of the three face-DCNNs are ResNet100, AlexNet, and VGG16, respectively. The ResNet100 face-DCNN was the pre-trained ArcFace model provided by the InsightFace package (labeled as “LResNet100E-IR,ArcFace@ms1m-refine-v2”, https://github.com/deepinsight/insightface/wiki/Model-Zoo#3-face-recognition-models), commonly known as the pre-trained ArcFace DCNN.

We trained the other two DCNNs also with the ArcFace loss function, one of the most effective loss functions for face recognition (74). To facilitate convergence, for each of these two DCNNs, we first trained it for 4 epochs using the softmax loss function, and then fine-tuned it for another 16 epochs using the ArcFace loss function. For the softmax loss function, we followed the steps of (74) to normalize the embedding vectors and weights.

We used the Adam optimizer for training (75), with an initial learning rate of 2^-10^. We used a procedure which we call “prestige” to choose the optimal learning rate. That is, for each epoch, we trained two replicas of the DCNN with different learning rates: one with the same learning rate as the previous epoch, and the other with half the previous learning rate. After both replicas had finished training, we only kept the one with a smaller loss. This procedure allows us to train our DCNNs with a satisfactory convergence speed. We also repeated the training of each network twice with different initializations, and only used the one with smaller loss.

We assessed the performance of the two DCNNs we trained using the Labeled Faces in the Wild (LFW) dataset (76). Specifically, we used the curated version of the dataset provided by the InsightFace package, which contained 6000 pairs of images.

We used a 10-fold cross-validation scheme to evaluate the prediction accuracy of our DCNNs. For each pair of images, we computed the similarity of their embedding vectors, and predicted whether they were the same identity or not based on a threshold. For each cross-validation fold, the threshold was chosen based only on training data.

The classification accuracy was 98.75% and 98.45% (chance accuracy: 50%) for the face AlexNet and face VGG16 (Figure S2), respectively, which were comparable to previous results based on DCNNs that had similar architectures (https://paperswithcode.com/sota/face-verification-on-labeled-faces-in-the).

### DCNN Models Trained for Object Recognition

We used pretrained AlexNet and VGG16 models provided by the torchvision package (https://pytorch.org/vision/stable/models.html), which were optimized for object recognition based on the ImageNet dataset. Although those networks were not specifically trained for face recognition, they were able to classify face identity to some extent, with an accuracy of 68.43% and 68.30% for the object AlexNet and object VGG 16, respectively (chance accuracy: 50%).

## Data Analysis

### Preprocessing

MRI data were preprocessed using fMRIPrep version 1.4.1 (77). T1-weighted images were corrected for intensity non-uniformity (78) and skullstripped using antsBrainExtraction.sh. High resolution cortical surfaces were reconstructed with FreeSurfer (79) using both T1-weighted and T2-weighted images, and then normalized to the fsaverage template based on sulcal curvature (80). Functional data were slice-time corrected using 3dTshift (81), motion corrected using MCFLIRT (82), distortion corrected using fieldmap estimate scans (one for each session), and then resampled to the fsaverage template based on boundary-based registration (83). After these steps, functional data were in alignment with the fsaverage template based on cortical folding patterns. The following confound variables were regressed out of the signal in each run: six motion parameters and their derivatives, global signal, framewise displacement (84), 6 principal components from a combined cerebrospinal fluid and white matter mask (aCompCor) (85), and up to second-order polynomial trends.

### Searchlight Hyperalignment

All three imaging datasets were hyperaligned (26–29) based on responses to the Grand Budapest Hotel (Figure S1). We first built a common model information space where patterns of fMRI responses to the Grand Budapest Hotel movie were aligned across subjects. Whole-cortex transformation matrices for each individual were calculated using a searchlight-based algorithm to project each participant’s cortical space into the common model information space. Transformation matrices were calculated for all 15 mm radius searchlights in each brain using an iterative procedure and Procrustes alignment, and then aggregated into a single matrix for each hemisphere. Transformation matrices for each participant were used to transform their Hyperface and dynamic localizer data into the common model space, so that all three imaging datasets were functionally aligned in the same common model information space.

### Searchlight Representational Similarity Analysis

We performed a searchlight representational similarity analysis to quantify the similarity between DCNN and neural representational geometries. Embeddings derived from the final fully-connected layer and the intermediate layers were used to build the representational dissimilarity matrix (RDM) of the DCNN networks. In detail, the stimulus face and its five key landmarks were automatically detected in each frame to create the aligned and cropped face image. The cropped face image was then fed into the DCNN as input, and passed through the layers. Each video clip comprised 120 frames, and the corresponding 120 vectors were averaged to obtain an average embedding vector for each clip. Neural responses to each stimulus of the video clip were averaged over the duration of 4 s in each cortical vertex after adjusting for a 5 s hemodynamic delay, and the RDM was built using pattern similarity across clips for each 10 mm searchlight in each participant. The Hyperface stimulus set included 707 stimuli. This resulted in 707 × 707 RDMs for the DCNN layers and for each searchlight per participant, with each element of the RDM reflecting the correlation distance (1 – Pearson’s *r*) between the response patterns elicited by the two stimuli in a pair (Figure 1). The neural RDMs were first averaged across participants in each searchlight, and Pearson’s *r* values were calculated to measure the similarity between the model and neural RDMs across all surface searchlights to generate the whole-brain correlation map. To assess the statistical significance of whole-brain correlation maps, we performed a permutation test by shuffling the labels of the 707 stimuli prior to recomputing the RDMs 1000 times for each intermediate layer and 5000 times for the final fully-connected layer. The false discovery rate (FDR) was controlled at *p* < 0.005 to obtain whole-brain FDR corrected maps. For run-by-run analysis, RSA was performed for each individual scanning run, and the correlation maps were averaged across runs.

### Correlations in the Face-Selective ROIs and the Noise Ceiling

The face-selectivity map was estimated using hyperaligned localizer data. We calculated the univariate contrast map of faces vs. objects for each participant using the hyperaligned localizer data in the common model information space, averaged these to get the group face-selective map (Figure S6), and applied a conservative threshold of *t* > 5 to obtain the face-selective regions. Individual ROIs including the occipital face area (OFA), the posterior and anterior fusiform face areas (aFFA and pFFA), the anterior temporal face area (ATL), the posterior and anterior superior temporal sulcus (pSTS and aSTS), and three inferior frontal face areas (superior, middle, and inferior: sIFG, mIFG, and iIFG) bilaterally were localized by drawing a disc of radius = 10 mm centered on the peak face-selective response (see also analysis with radius = 15 mm and 20 mm in Figure S10 & S11). Mean correlation coefficients were calculated for searchlights with centers within face-selective areas and non-face-selective areas for each layer of DCNNs. The correlation coefficient of each face-selective ROI was the value for the searchlight centered on the peak of face-selective response. Standard errors of the mean were calculated by bootstrapping the stimuli 1000 times for each intermediate layer and 5000 times for the final fully-connected layer. Statistical significance was assessed by permutation tests randomizing the stimulus labels 1000 times for each intermediate layer and 5000 times for the fully-connected layer.

The noise ceiling provides an estimate of the maximum possible correlation with the neural RDM predicted by the unknown true model (86). Because we averaged individuals’ RDMs before RSA analysis, the noise ceiling was estimated by calculating Cronbach’s alpha using neural RDMs across participants (87). Cronbach’s alpha was used to describe the reliability of the neural RDMs across participants in each searchlight. To obtain noise ceilings for the face-selective areas, non-face-selective areas, and across the whole brain, mean alphas were calculated by averaging across vertices (corresponding to searchlight centers) in these regions. For the run-by-run analysis, the noise ceiling was estimated for each individual scanning run first and was averaged across runs to get the estimation of the overall noise ceiling.

### Reweighting Features Prior to RSA

RSA has the strong assumption that all features contribute equally to generate an RDM (e.g. all cortical vertices in a searchlight are equally important when computing pattern similarity between two conditions) (40, 41). We tested whether relaxing this assumption might yield larger DCNN-neural correlations. We developed a novel approach that best matched the DCNN features to the brain responses before performing RSA. First, the DCNN features were matched to the brain responses by performing singular value decomposition (SVD) on the covariance matrix between the DCNN features and brain features. This step corresponds to bringing the DCNN features into a space with reduced dimensionality *N* that best matches brain responses. Second, ridge regression was used to predict responses for each brain feature (vertex) from the reduced-dimension DCNN features. Third, the fitted ridge model was used to generate predictions of the brain responses from DCNN features for left-out data. This procedure ultimately yields a reweighted subspace of the original DCNN feature space that best predicts brain features. Finally, these predicted features were used to generate RDMs, which were then analyzed as reweighted DCNN RDMs.

This process was performed using nested cross-validation. The 12 scanning runs were separated into two sets (11 training runs and one test run) for both the brain and DCNN (ArcFace) responses. The training runs were used to estimate the transformation for the shared components and the ridge regression parameters. The hyperparameters (number of dimensions *N* and ridge regularization parameter α) were chosen based on a nested leave-one-run-out loop within the 11 training runs. We performed a grid search on *N* and α by testing 18 evenly distributed values from 5 to 90 for *N*, and 29 evenly distributed values from 10*^-7^* to 10*^7^* on a logarithmic scale for α. The regression model was trained to yield the best *R^2^*. The best model was then used to generate RDMs for the left-out run. This analysis was performed within each searchlight.

### Cross-subject Identity Decoding

The cross-subject identity decoding analysis was done as a binary classification task with a simple one-nearest-neighbor classifier across all searchlights (10 mm radius). We performed the analysis using a split-half cross-validation scheme. That is, we divided the subjects into two groups (training and test). For each face and each group, we computed an average response pattern across subjects in the group. We assessed whether the average pattern of a face for the training group is more similar to that of the same face for the test group compared to a different face (2-alternative forced choice; chance accuracy = 50%). We repeated this for all pairs of faces and averaged the accuracy. Because there are many different ways to split the subjects into two groups, we also repeated the split-half procedure 100 times and averaged the accuracy across repetitions. We also performed a similar analysis using leave-one-subject-out cross-validation instead of split-half cross-validation. Each time, we computed the average response patterns of 20 subjects and compared with the left-out test subject.

Note that the RSA analysis is based on the average of all 21 subjects rather than half of the subjects or a single subject, and the data quality is superior to the average patterns used in this classification analysis.

### Behavioral Arrangement Task RDMs and Noise Ceilings

Coordinates at the center point of the thumbnails were used to build behavioral RDMs for stimuli in each scan run for each participant. Each element of the behavioral RDMs reflected the Euclidean distance between the placements for a given pair of stimuli. Individual behavioral arrangement task RDMs were averaged across participants before further analysis. Because we averaged individual behavioral RDMs in each run before further analysis, the noise ceiling for each run was estimated using Cronbach’s alpha across participants and averaged across runs.

### Variance Partitioning Analysis

Variance partitioning analysis based on multiple linear regression was used to quantify the unique contributions of each model taking into consideration the contribution of other models. In detail, this analysis was done to tease apart the contribution of face- and object-trained DCNNs in explaining variance of the behavioral arrangement task RDM, and to separate the relative contributions of face- and object-trained DCNNs, as well as the behavioral arrangement task performance in explaining variance of the neural RDM for a given face-selective region. In the first analysis (comparing behavioral RDMs and two types of DCNN RDMs), the off-diagonal elements of the behavioral RDM were assigned as the dependent variable, and the off-diagonal elements of the two DCNN models were assigned as independent variables (predictors). In the second analysis (comparing neural, behavioral, and two types of DCNN RDMs), the dependent variable was the off-diagonal elements of the mean RDM across vertices in face-selective cortex, and the independent variables were the off-diagonal elements in the two DCNN models and the behavioral RDM. Both analyses were performed within runs and the variance was averaged across runs as the final result. To obtain unique and shared variance for each model, in the former analysis, three multiple regression analyses were run in total. The three analyses included the full model that had both DCNNs as predictors and two reduced models that contained an individual DCNN as the predictor. For the latter analysis, seven multiple regression analyses were run including one full model that had behavioral and two DCNN models as predictors, as well as six reduced models that had either combinations of two models from the three (behavioral, two DCNNs) or one individual model alone as the predictor. By comparing the adjusted explained variance (adjusted *R*^2^) of the full model and the reduced models, variance that was explained by each model independently could be inferred (88, 89). The variance partitioning analysis was conducted using the “vegan” package of R (https://cran.r-project.org/web/packages/vegan/vegan.pdf).

RSA between the behavioral arrangement task RDMs and RDMs of DCNNs were carried out for each run and correlations were averaged across runs. “Best layers” for the behavioral arrangement task were layers that had the strongest correlations with the behavioral RDMs, including _plus45 of ArcFace, pool5 of face AlexNet, block5_conv2 of face VGG16, fc2 of object AlexNet, and fc2 of object VGG16. In the run-by-run RSA analysis between neural RDMs and RDMs of DCNNs, “best layers” were_plus42 of ArcFace, conv1 of face AlexNet, block2_pool of face VGG16, conv2 of object AlexNet, and block3_pool of object VGG16. These layers were very similar with the “best layers” in the RSA analysis using all 707 stimuli (_plus42 of ArcFace, conv1 of face AlexNet, block2_conv1 of face VGG16, conv2 of object AlexNet, and block3_conv3 of object VGG16).

### Multidimensional Scaling

Multidimensional scaling (MDS) was used to visualize the representational geometry of face stimuli for different DCNNs, behavioral arrangements, and neural ROIs. Pairwise correlation distance matrices were computed for stimuli pairs in each of the spaces in a run-by-run manner. Metric MDS was used to project the stimuli onto a 2-dimensional space and these locations were color-coded based on the behavioral ratings. MDS was also used to visualize representational geometry of the 18 face-selective ROIs (Figure S5). Pairwise correlation distance matrices were computed for face ROI pairs based on the 707-by-707 RDM of each face ROI in a second-order RSA manner.

## Acknowledgements

We thank the authors of the InsightFace package for making their models and training data freely available for non-commercial research use. This work was supported by NSF grants 1607845 (J.V.H) and 1835200 (M.I.G).

## Author contributions

Conceptualization, G.J., M.F., J.V.H., and M.I.G.; Methodology, G.J., M.F., J.V.H., and M.I.G.; Software, G.J., M.F., and S.A.N.; Formal Analysis, G.J. and M.F.; Investigation, G.J., M.F., and M.V.d.O.C.; Resources, J.V.H. and M.I.G.; Data Curation, G.J., M.F., and M.V.d.O.C.; Writing – Original Draft, G.J., M.F., J.V.H., and M.I.G.; Writing – Review & Editing, G.J., M.F., M.V.d.O.C., S.A.N., J.V.H., and M.I.G.; Visualization, G.J. and M.F.; Supervision, J.V.H. and M.I.G.; Funding Acquisition, J.V.H. and M.I.G.

## Data and materials availability

All data needed to evaluate the conclusions in the paper are present in the paper and/or the Supplementary Materials. Further data and codes can be downloaded from https://github.com/GUO-Jiahui/face_DCNN. Additional data and materials that support the findings of this study are available on request from the corresponding authors G.J., M.F., and M.I.G.

## Supplementary Materials

**Figure S1.**
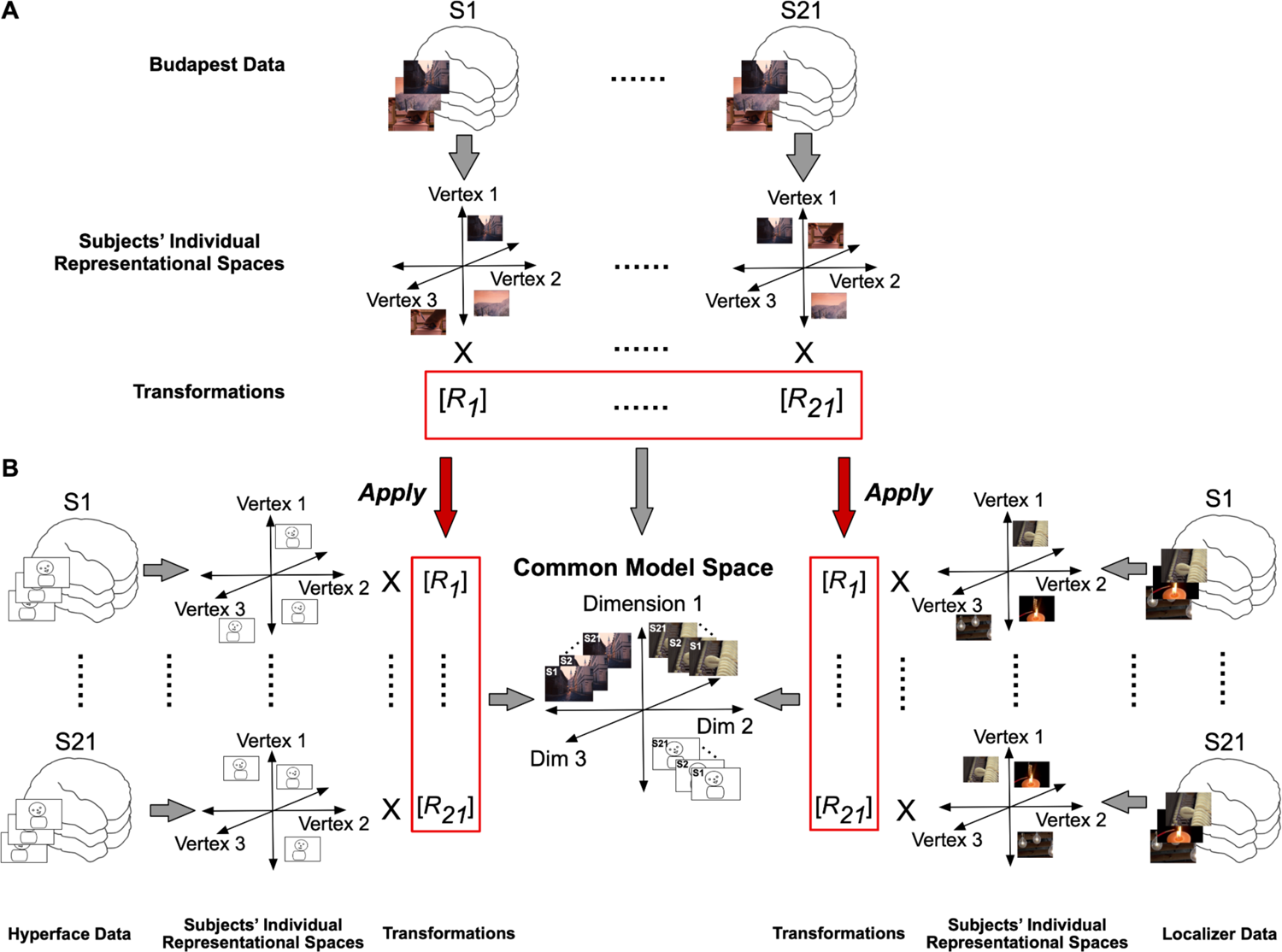
Schematic of the hyperalignment procedure. **A.** Transformation matrices were calculated by hyperaligning each participant’s responses to the Grand Budapest Hotel movie to the common model space. **B.** The transformation matrix derived from the Grand Budapest Hotel movie for each participant was then applied to the Hyperface data. **C.** The transformation matrix derived from the Grand Budapest Hotel movie for each participant was also applied to the localizer runs. Thus, after hyperalignment, the Budapest data, the Hyperface data, and the localizer data were all functionally aligned to the common model space.

**Figure S2.**
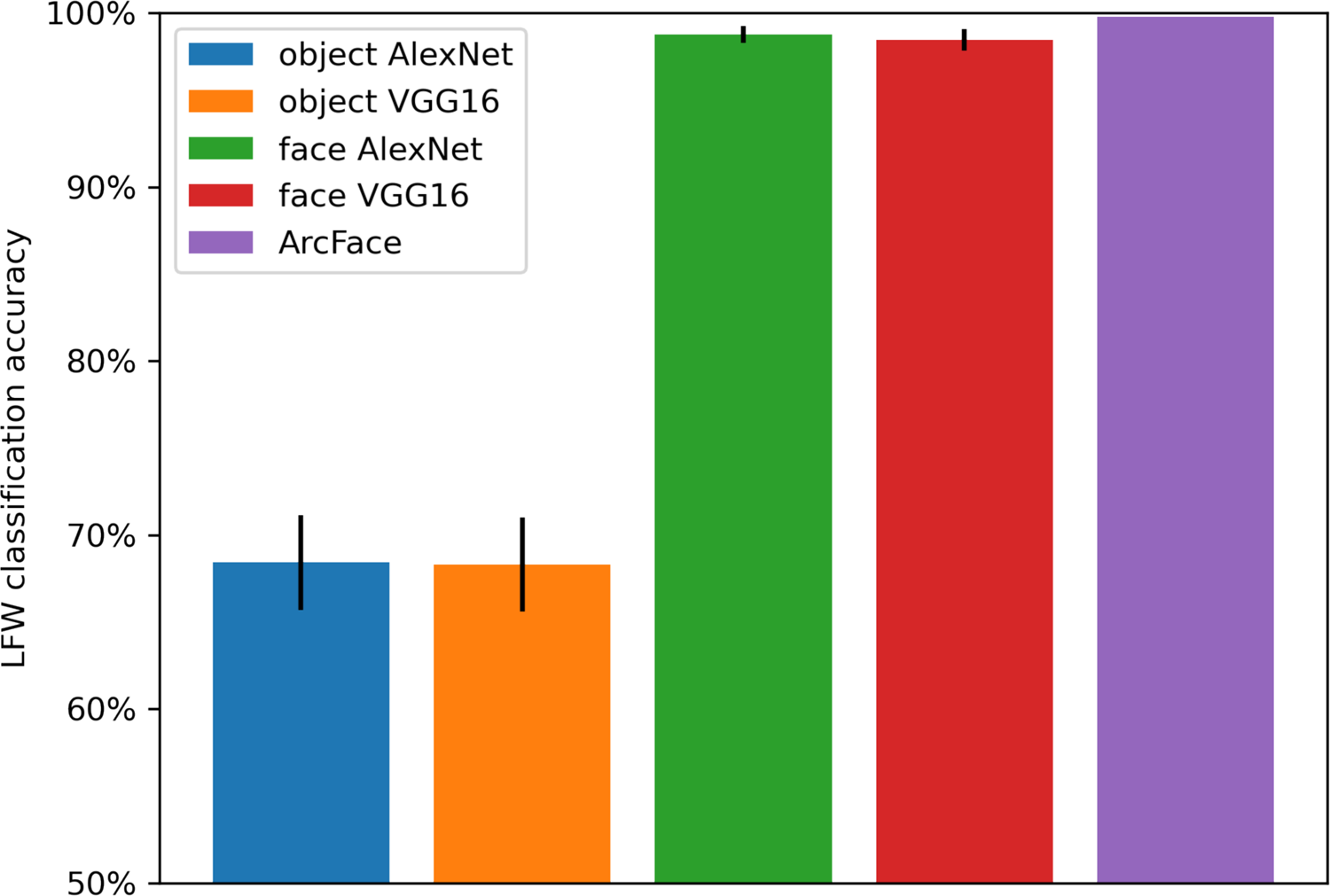
DCNN performance on face recognition. Using the Labeled Faces in the Wild (LFW) dataset, we assessed the face recognition performance of the DCNNs used in our analysis. The benchmarking task was to tell if two faces were the same person or not based on the similarity of their embeddings, and thus the chance accuracy was 50%. The dataset was divided into 10 folds and a leave-one-out cross-validation was used to determine the threshold (i.e. the similarity level to determine if two faces are the same or not). The accuracy was 68.43% and 68.30% for the object AlexNet and object VGG16, respectively, and 98.75% and 98.45% for the face AlexNet and face VGG16, respectively. The accuracy was 99.77% for the pre-trained ArcFace, which was provided by the InsightFace package. Error bars are the standard deviation of the accuracy across 10 cross-validation folds.

**Figure S3.**
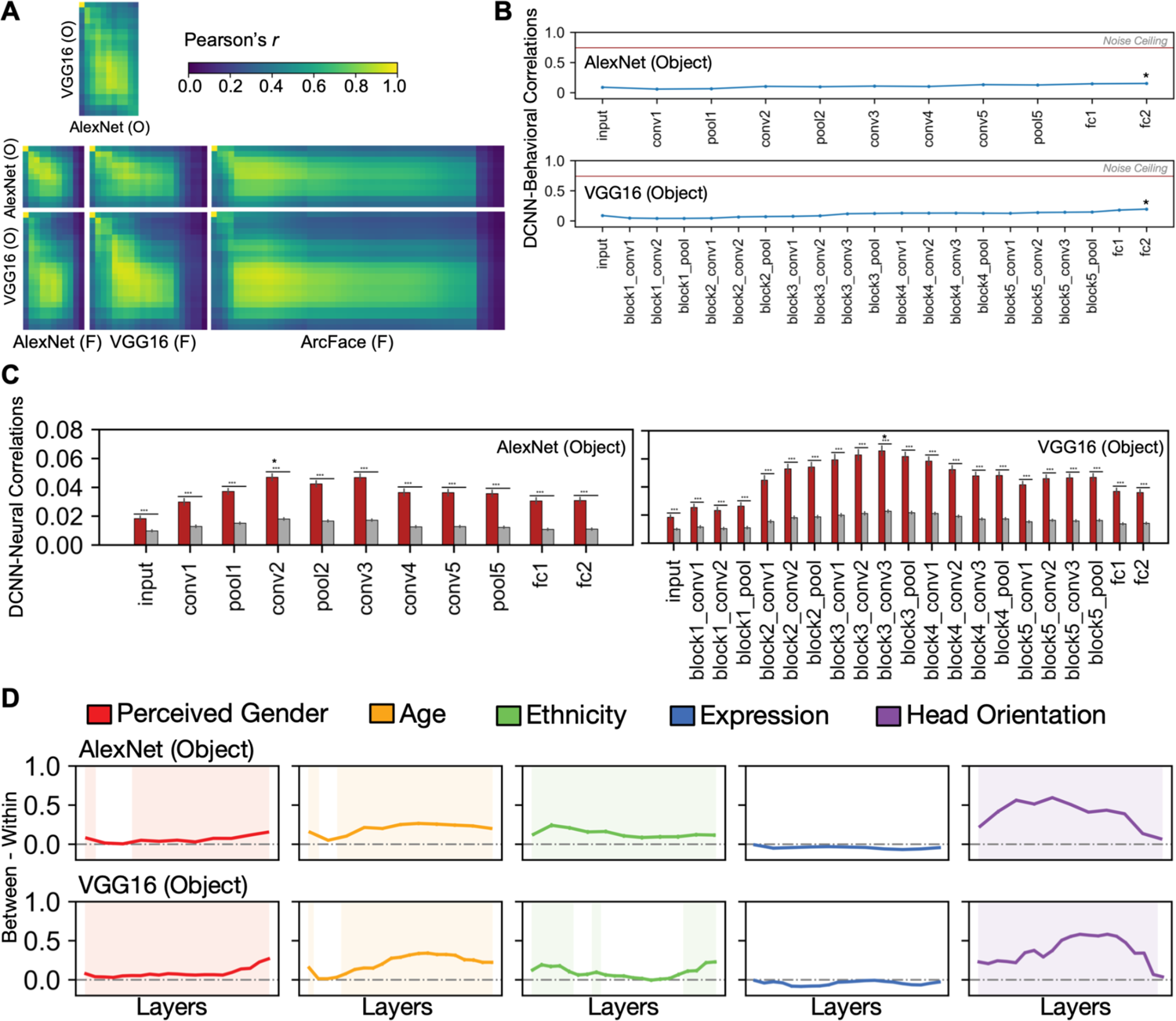
Object-DCNN representations. **A.** Correlations in each pair of layers in the two object-trained AlexNet and VGG16, and across the six face-(ArcFace, AlexNet, and VGG16) and object-DCNN pairs. **B.** Mean correlations across participants and runs between the behavioral RDM and DCNN RDM in each layer in the two object-DCNNs. The star marks the layer that has the highest correlation with the behavioral task in each DCNN. The red horizontal line in each subplot represents the mean noise ceiling of the behavioral arrangement task across runs. **C.** Average correlations for face-selective regions and non-face-selective regions for each layer in the two object-DCNNs. Regions, significance, and the color code were defined the same as in Figure 3C. Stars indicate the layers that had the largest correlations. **D.** Difference in the between- and within-group distance of perceived gender (red), age (orange), ethnicity (green), expression (blue), and head orientation (purple) in representational geometries of the behavioral arrangement task, each layer of the two object-trained DCNNs (AlexNet, VGG16). These differences were calculated within each run and then averaged across runs. Shaded layers and ROIs show significant differences in the between-versus within-group test (*p* < 0.05, permutation test, one-tail). Significance of the difference was estimated based on a random permutation test randomizing the stimulus labels. Error bars represent one standard error of the mean estimated by bootstrap resampling stimuli. *** *p* < 0.001.

**Figure S4.**
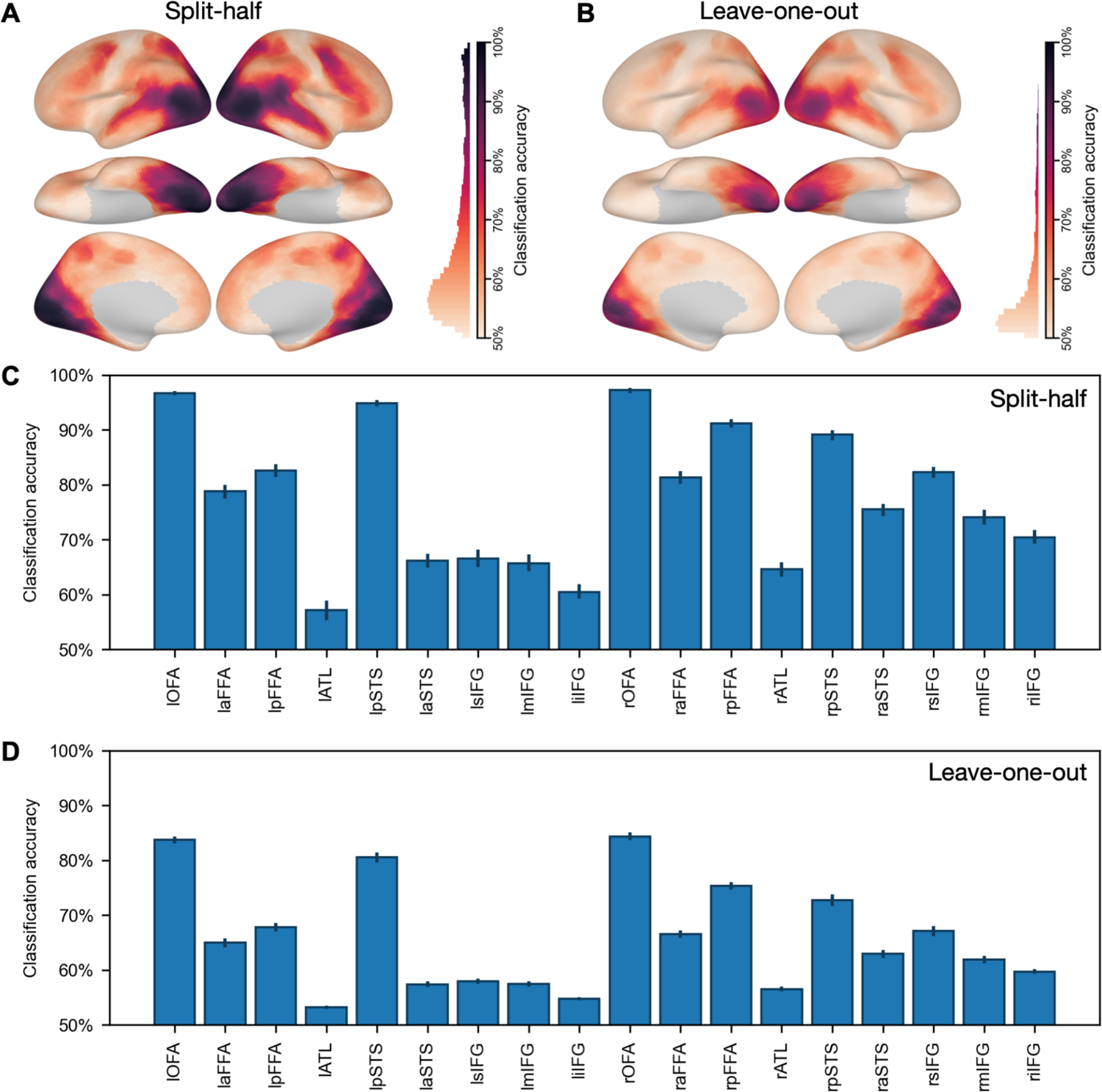
Between-subject identity decoding accuracies. A. Searchlight analysis of between-subject identity classification. This classification analysis was a binary classification task with a simple one-nearest-neighbor correlation-based classifier. We used a searchlight radius of 10 mm, similar to our other analyses. We used a split-half cross-validation scheme. That is, we divided the subjects into two groups (training and test). The classification task is binary classification (2-alternative forced choice). For each face and each group, we computed an average response pattern across subjects in the group. Each time, we assessed whether the test group’s average pattern of a certain face was more similar to the training group’s pattern to the same face than to a different face (chance accuracy = 50%). We repeated this procedure for each pair of faces and averaged the accuracy across face pairs. Because there are multiple ways to split subjects into groups, we also repeated the split-half procedure 100 times and averaged the accuracy across repetitions. Classification accuracy was high for visual and face-selective regions. B. Similar analysis based on a leave-one-subject-out cross-validation scheme. Each time, we computed the average response patterns of 20 subjects and compared these patterns with those of the left-out test subject. The accuracies were lower compared with A, because the data from only a single subject was used as test data, but nonetheless clearly significant. Note that even in the split-half condition, there were only 10 or 11 subjects per group. Our RSA analysis was based on the average across all 21 subjects, and thus the average patterns have higher quality than those used in A. C. Split-half classification accuracy in face-selective regions. D. Leave-one-subject-out classification accuracy in face-selective regions.

**Figure S5.**
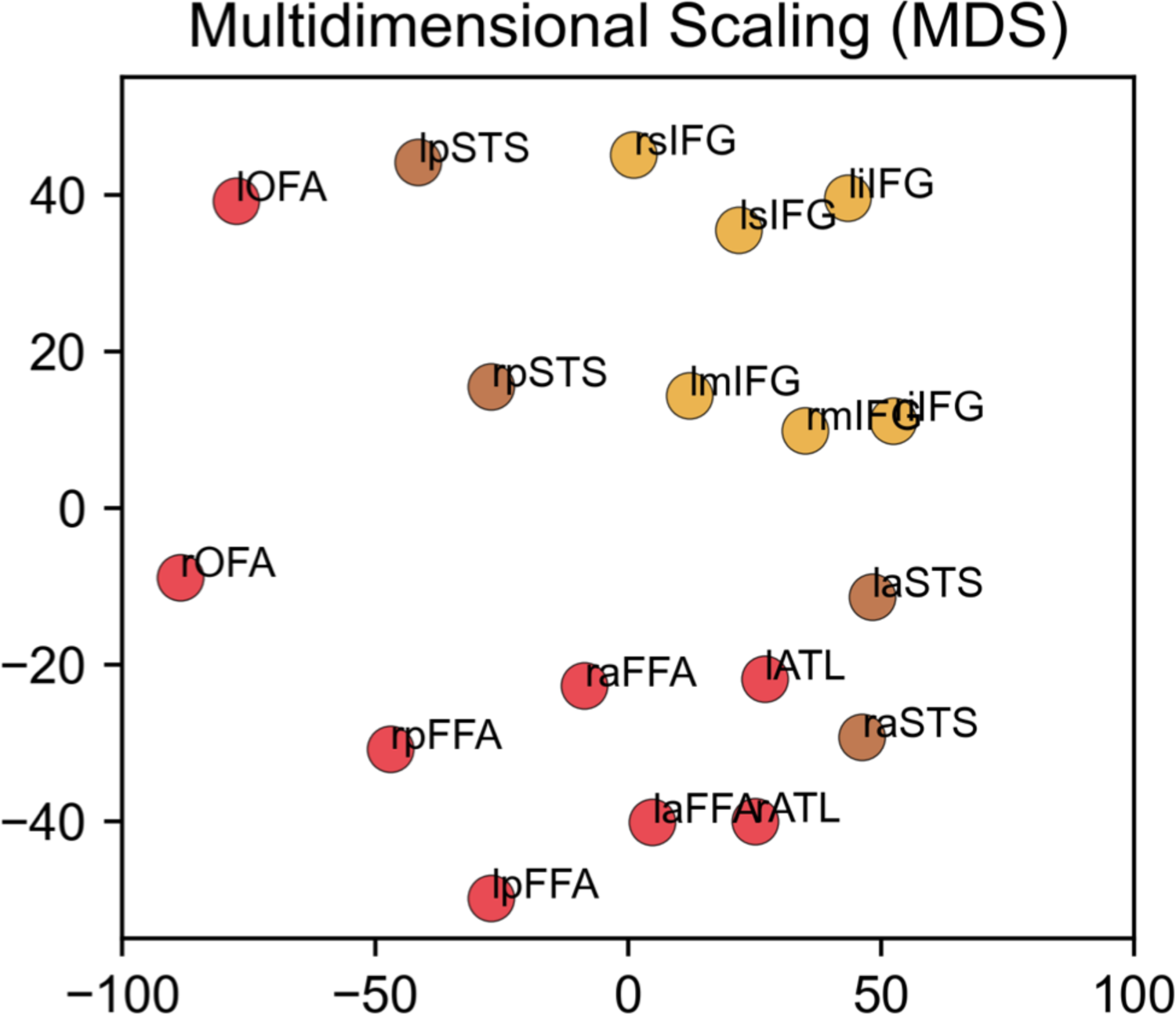
MDS plot of distances between RDMs in 18 bilateral face-selective ROIs. Each dot represents one face-selective ROI (10 mm radius with the peak at the center). These face ROIs were localized using the faces-vs-objects contrast with the dynamic localizer. Red markers denote ROIs in ventral temporal cortex, orange markers denote ROIs in lateral temporal cortex, and yellow markers denote ROIs in frontal cortex. The clustering of ROIs follows the anatomical and functional hierarchy of the face-processing network.

**Figure S6.**
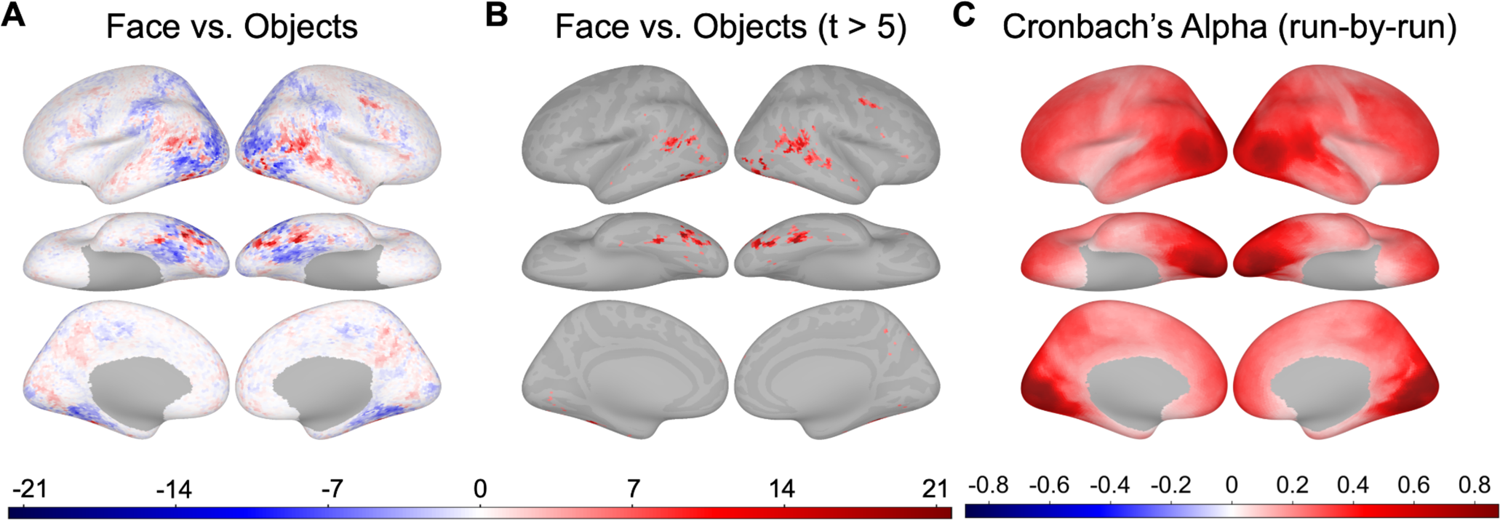
Contrast maps and noise ceilings. **A.** The whole-brain faces-vs-objects contrast *t*-map was calculated for each individual using hyperaligned dynamic localizer data. Then the *t*-maps were averaged across participants. The darker the red color was, the stronger the responses were to faces. **B.** The face-selective regions thresholded at *t* > 5. **C.** Whole-brain Conbach’s alpha map (run-wise). Reliabilities of neural RDMs across participants were calculated for each searchlight for each run, and the mean Cronbach’s alpha map was derived by averaging maps across runs.

**Figure S7.**
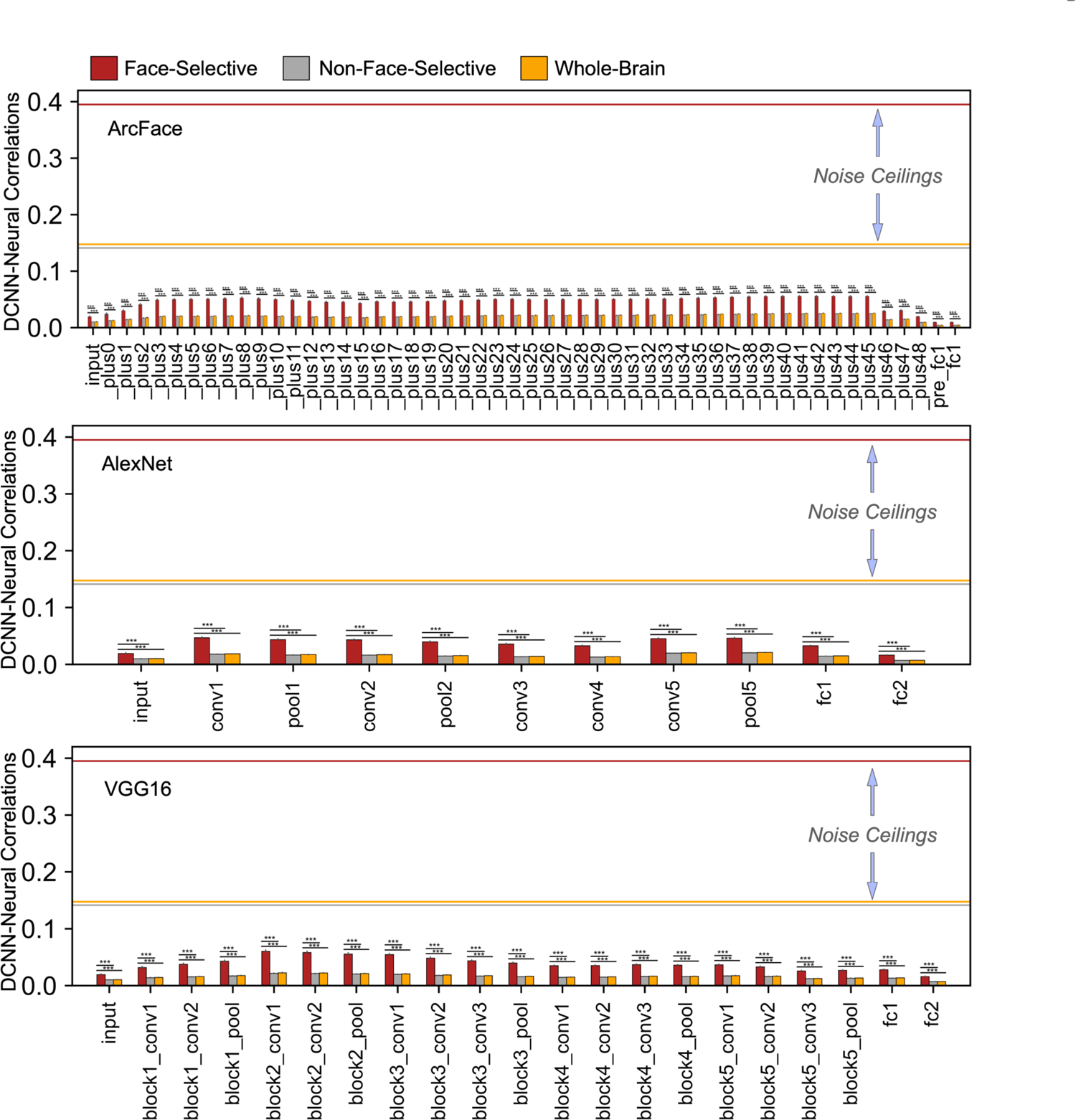
RSA results in layers of three face-DCNNs with noise ceilings. Bar plots of the mean DCNN-neural correlations in face-selective, non-face-selective, and whole-brain regions in layers of the three face-DCNNs with noise ceilings. Horizontal lines in each panel represent the mean noise ceiling in face-selective, non-face-selective, and whole-brain searchlights. These lines are coded with the same colors in the legends. In all three panels, error bars represent one standard error of the mean estimated by bootstrap resampling stimuli. Significance of the difference was estimated based on a permutation test randomizing the stimulus labels. *** *p* < 0.001.

**Figure S8.**
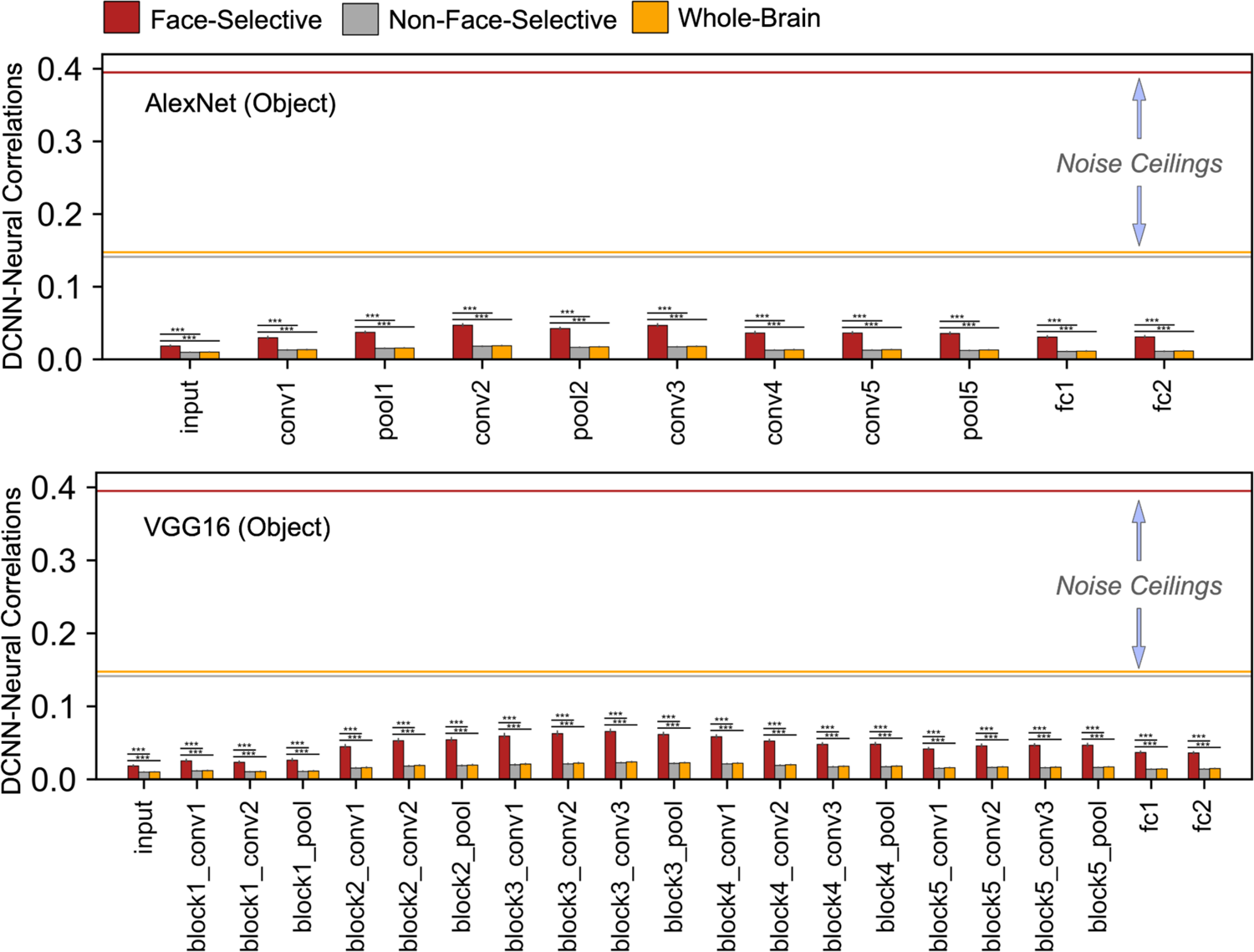
RSA results in layers of two object-DCNNs with noise ceilings. Bar plots of the mean DCNN-neural correlations in face-selective, non-face-selective, and whole-brain regions in layers of the two object-DCNNs with noise ceilings. Horizontal lines in each panel represent the mean noise ceiling in face-selective areas, non-face-selective areas, and across the whole brain. These lines are color coded according to the legend at top. In panels A, B, and C, the error bars indicate standard error of the mean estimated by bootstrap resampling stimuli. Significance of the difference was assessed based on a permutation test randomizing the stimulus labels. *** *p* < 0.001.

**Figure S9.**
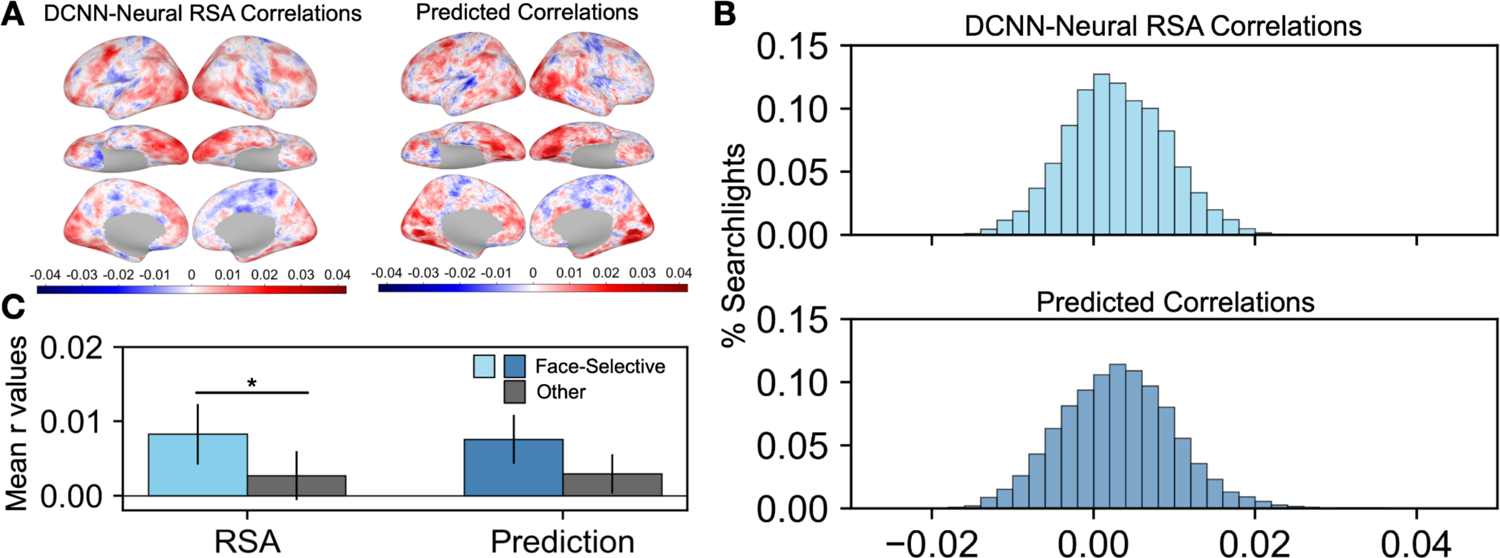
RSA and prediction analysis. **A.** Left panel shows the whole-brain ArcFace-neural RSA map (run-wise). Right panel shows the whole-brain prediction correlation map. Because prediction analysis was done in a leave-one-run-out cross-validation procedure (see Material & Methods for details), the RSA was recalculated using a run-wise procedure for comparison. In detail, RDMs were computed for each run in each searchlight, and the resulting correlations were averaged across runs to generate the map in the left panel. **B.** Histograms of ArcFace-neural RSA correlations calculated using data in each run and averaged across all twelve runs are plotted at bin size of 0.002 in the upper panel. Histograms of correlations between predicted and original RDMs are plotted at the same bin size in the lower panel. **C.** Average correlations of face-selective regions (*t* > 5) and non-face-selective regions (*t* <= 5) using RSA and prediction maps. The error bars indicate standard error of the mean estimated by bootstrap resampling of the stimuli (5000 bootstraps). Significance of the difference between the two bars was assessed using a permutation test randomizing the stimulus labels (5000 permutations). * *p* < 0.05.

**Figure S10.**
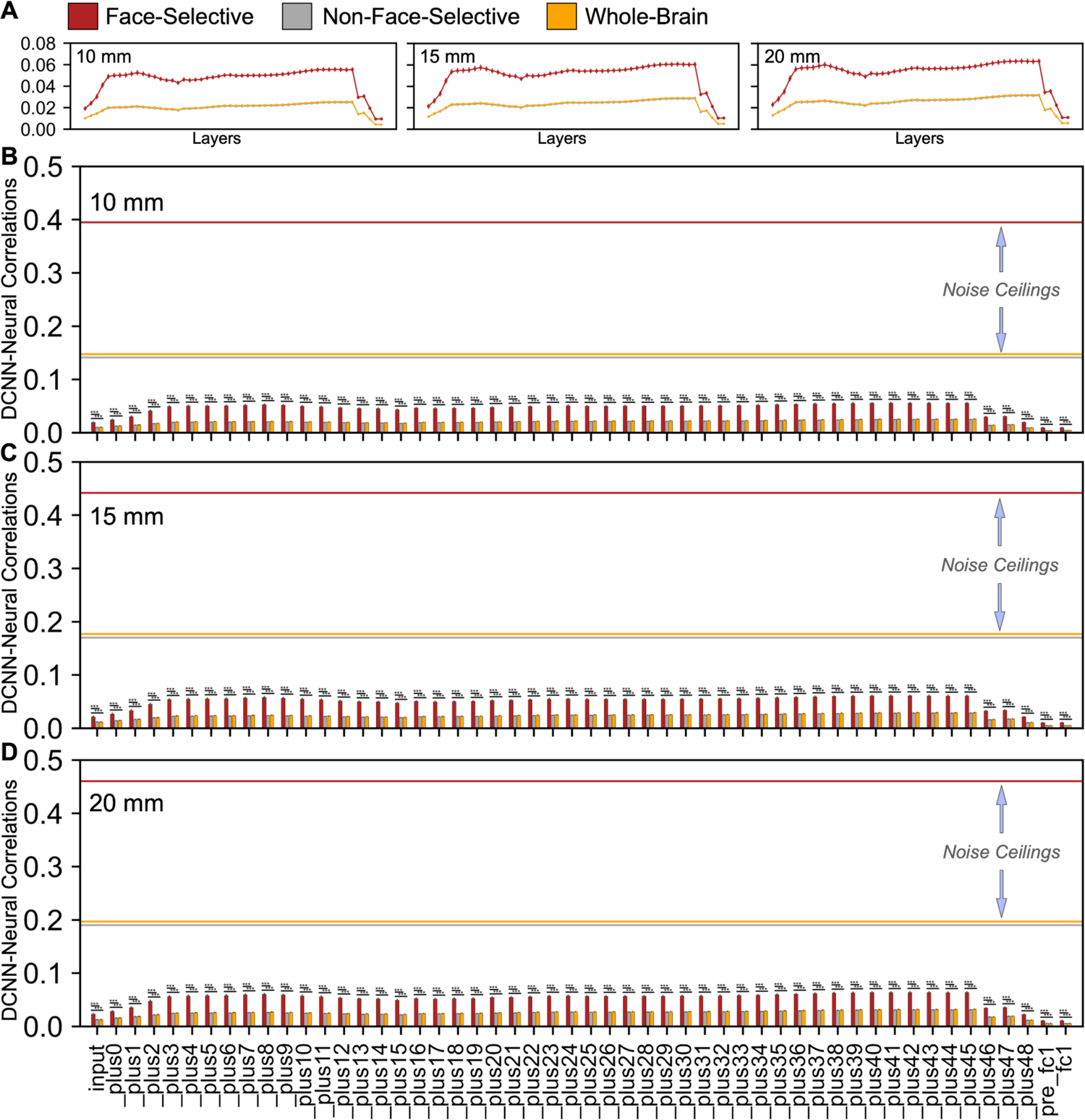
RSA results in layers of ArcFace with different searchlight sizes. **A.** Line plots of the mean DCNN–neural correlations in face-selective regions (red), non-face-selective regions (gray), and across the whole brain (orange) for each layer of ArcFace with a searchlight radius of 10 mm, 15 mm, and 20 mm (note that the gray and orange lines are largely overlapped). **B, C, & D.** Bar plots of the same values as in panel A (mean DCNN-neural correlations in face-selective, non-face-selective, and whole-brain regions in layers of ArcFace) with noise ceilings. Horizontal lines in each panel represent the mean noise ceiling in face-selective, non-face-selective, and whole-brain searchlights. These lines are coded with the same colors in the legends. In all four panels, error bars represent one standard error of the mean estimated by bootstrap resampling stimuli. Significance of the difference was estimated based on a permutation test randomizing the stimulus labels. *** *p* < 0.001. Larger searchlight sizes slightly increased DCNN-neural correlations, but the overall correlations remained weak.

**Figure S11.**
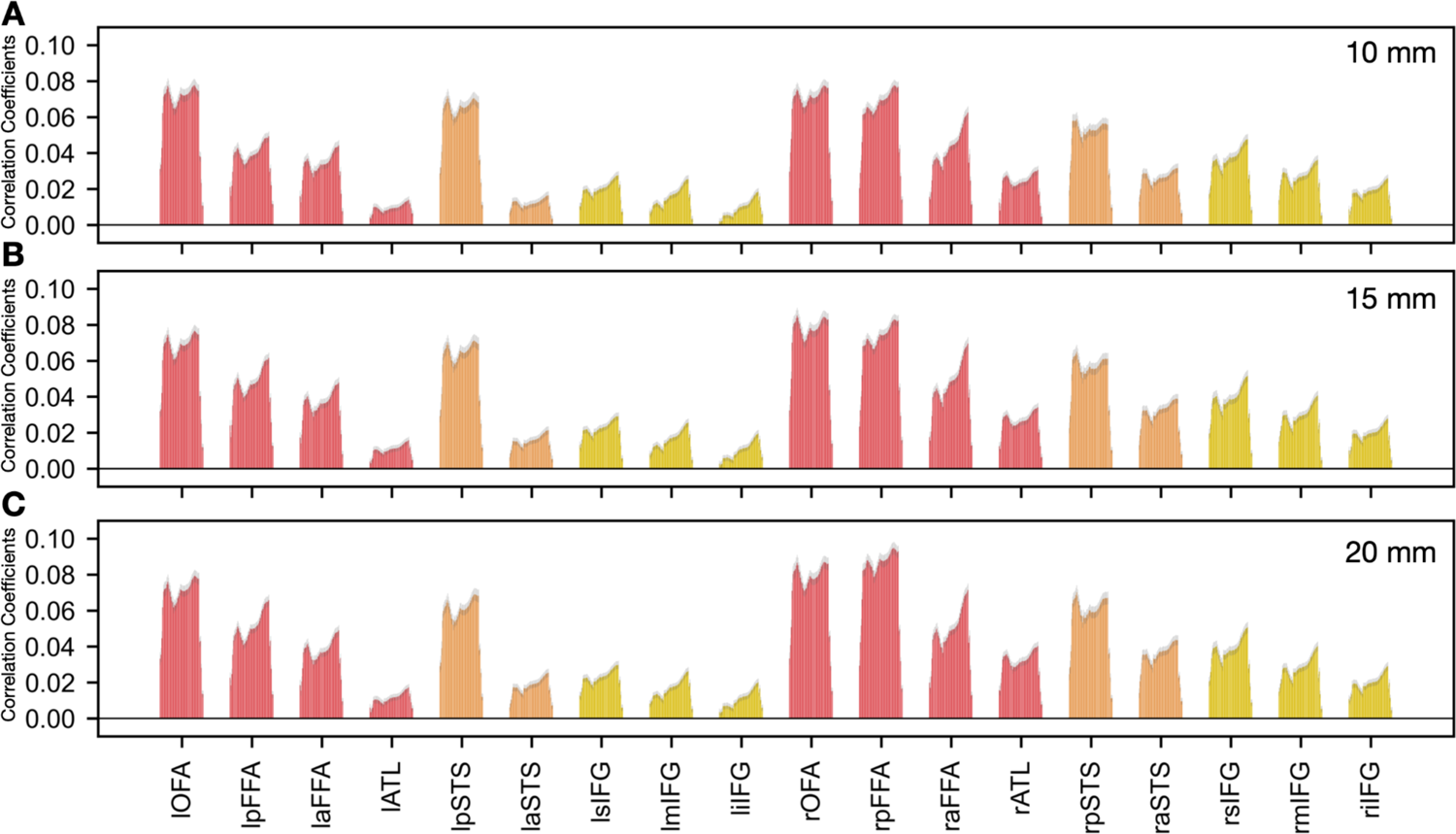
RSA results in individual face-selective ROIs with different ROI sizes. **A, B, & C.** RSA correlations for each layer of ArcFace in individual face-selective ROIs with ROI size of r = 10 mm, 15 mm, and 20 mm. Error bars indicate standard error of the mean estimated by bootstrap resampling the stimuli (1000 bootstraps).

**Figure S12.**
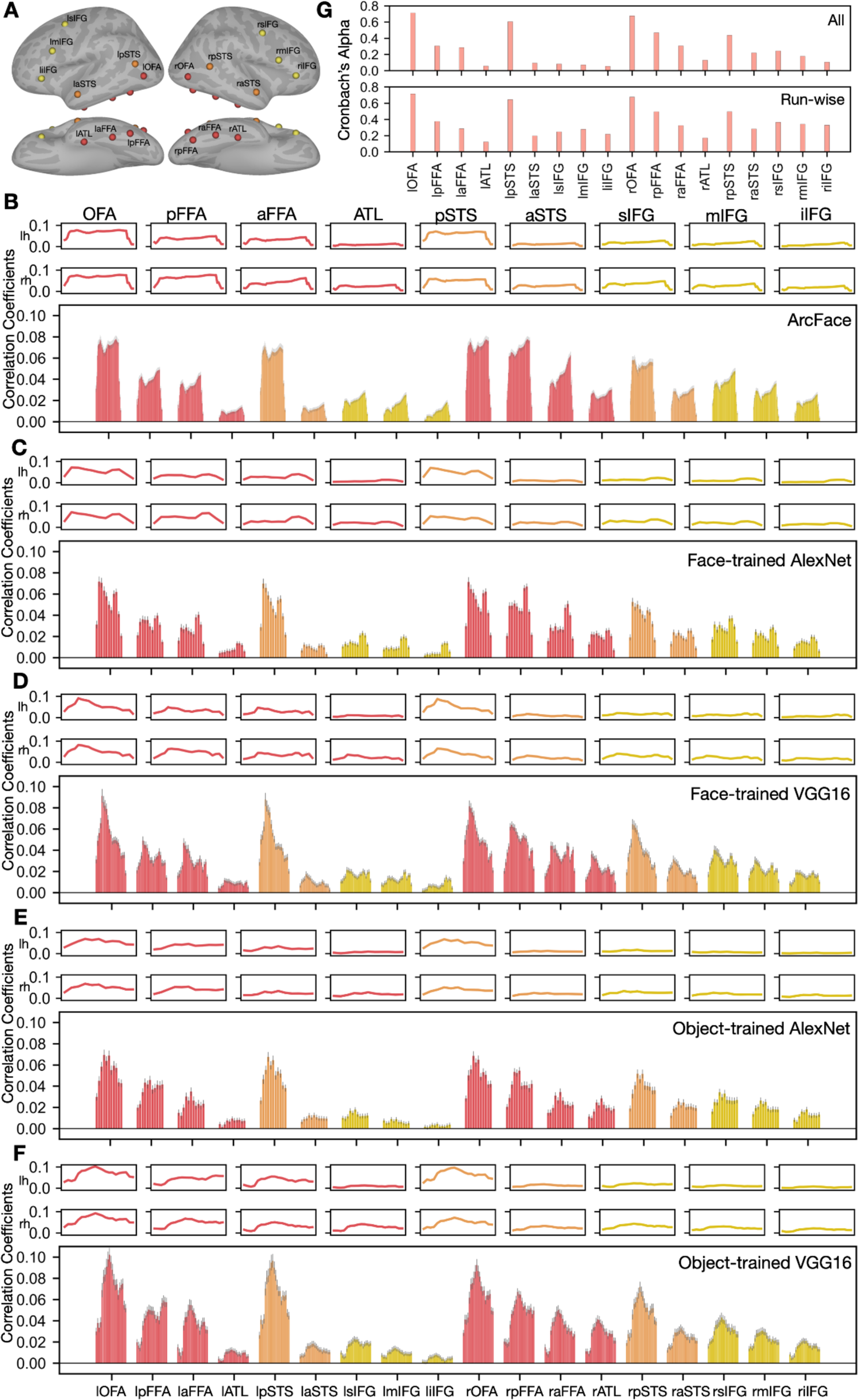
RSA results and noise ceilings in individual face-selective ROIs. **A.** Peak coordinates of 18 bilateral face-selective ROIs across both hemispheres. Red markers denote ROIs in ventral temporal cortex, orange markers denote ROIs in lateral temporal cortex, and yellow markers denote ROIs in frontal cortex. **B, C, D, E, & F.** RSA correlations for each layer of ArcFace, AlexNet (face-trained), VGG16 (face-trained), AlexNet (object-trained), VGG16 (object-trained) respectively in individual face-selective ROIs. Error bars indicate standard error of the mean estimated by bootstrap resampling the stimuli (1000 bootstraps). The line plots at the top of each panel match the values from the bar plot below. The line plots are included to more clearly display the profile of correlations across DCNN layers. **G.** Noise ceilings (Cronbach’s alphas) in each face-selective ROI. Noise ceilings were calculated using all stimuli at once (All, upper panel), and within each run and averaged across runs (Run-wise, lower panel). The DCNN-neural correlations were much lower than the noise ceiling of that ROI across all layers.

**Figure S13.**
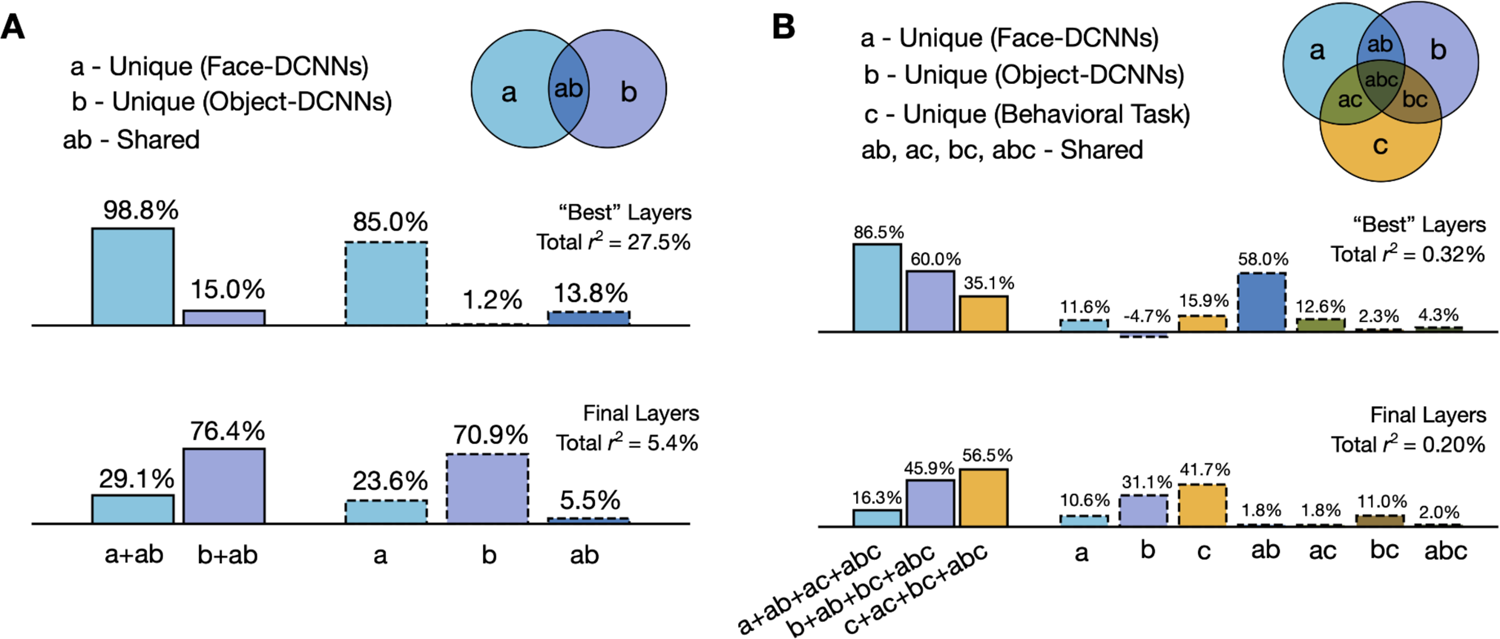
Variance partitioning analysis. **A.** Results of the variance partitioning analysis on the RDMs of face-DCNNs (light blue) and the object-DCNNs (purple) explaining variance of the behavioral task representations. Variance noted as unique or shared (numbers above the bars) are percentages of the total variance explained by both models combined (total *r*^2^). The first two bars (solid edges) show the total variance that each DCNN category (face or object) explained, and the rest of the bars (dotted edges) show the unique and shared variance that each DCNN category explained. In the upper panel, face- and object-DCNN RDMs were the mean of the RDMs in layers with the highest correlations with the behavioral RDM (“best” layers, marked with stars in panel B. _plus45, pool5, block5_conv2, fc2, and fc2 in ArcFace, face AlexNet, face VGG16, object AlexNet, and object VGG16 accordingly). In the lower panel, face- and object-DCNN RDMs were the mean of the RDMs in the final layer. **B.** Results of the variance partitioning analysis on the RDMs of face-DCNNs (light blue), object-DCNNs (purple), and behavioral task (yellow) explaining variance of the neural representations in face-selective areas (*t* > 5). Variance noted as unique or shared (numbers above the bars) are percentages of the total variance explained by all three models combined (total *r*^2^). The first three bars (solid edges) stand for the total variance that DCNNs and the behavioral task explained, and the rest of bars (dotted edges) stand for the unique and shared variance that each DCNN category and the behavioral task explained. In the upper panel, face- and object-DCNN RDMs were the mean of the RDMs in layers that had the highest correlations with the neural RDM in the run-by-run analysis (“best” layers, _plus42, conv1, block2_pool, conv2, block3_pool in ArcFace, face-trained AlexNet, face-trained VGG16, object-trained AlexNet, and object-trained VGG16 accordingly). In the lower panel, face- and object-DCNN RDMs were the mean of the RDMs in the final layer.

